# Plasma membrane damage limits replicative lifespan in yeast and induces premature senescence in human fibroblasts

**DOI:** 10.1101/2021.03.26.437120

**Authors:** Kojiro Suda, Yohsuke Moriyama, Yumiko Masukagami, Nurhanani Binti Razali, Yatzu Chiu, Koutarou Nishimura, Hunter Barbee, Hiroshi Takase, Shinju Sugiyama, Yoshikatsu Sato, Tetsuya Higashiyama, Yoshikazu Johmura, Makoto Nakanishi, Keiko Kono

## Abstract

Plasma membrane damage (PMD) occurs in all cell types due to environmental perturbation and cell-autonomous activities. However, cellular outcomes of PMD remain largely unknown except for recovery or death. Here, using budding yeast and normal human fibroblasts, we show that cellular senescence, irreversible cell cycle arrest contributing to organismal aging, is the long-term outcome of PMD. To identify the genes essential for PMD response, we developed a simple PMD-damaging assay using a detergent and performed a systematic yeast genome-wide screen. The screen identified 48 genes. The top hits in the screen are the endosomal sorting complexes required for transport (ESCRT) genes, encoding the well-described plasma membrane repair proteins in eukaryotes. Unexpectedly, the replicative lifespan regulator genes are enriched in our 48 hits. This finding suggests a close genetic association between the PMD response and the replicative lifespan regulations. Indeed, we show that PMD limits the replicative lifespan in budding yeast; the ESCRT activator AAA-ATPase *VPS4*-overexpression extends it. These results suggest that PMD limits replicative lifespan in budding yeast. Moreover, in normal human fibroblasts, we find that PMD induces premature senescence via the Ca^2+^-p53 axis but not the major senescence pathway, ATM/ATR pathway. Consistent with the results in yeast, transient overexpression of ESCRT-III, CHMP4B, suppressed the PMD-dependent senescence in normal human fibroblasts. Our study proposes that PMD limits cellular lifespan in two different eukaryotic cell types and highlights an underappreciated but ubiquitous senescent cell subtype, namely PMD-dependent senescent cells.

## Introduction

Cells experience a variety of perturbations on the plasma membrane ranging from physical attack to pathogen invasion. The plasma membrane is also damaged by physiological activities including muscle contraction (1, 2, 3). There is a growing appreciation that failed plasma membrane damage (PMD) response causes various diseases. For example, mutations in the PMD-repair protein dysferlin can lead to one form of muscular dystrophy (4); mutations in the PMD response factor lipid scramblase TMEM16F cause Scott syndrome (5, 6).

The PMD response is considered to be classified into two simple outcomes of recovery or death. Regarding the recovery response, plasma membrane repair mechanisms, including Ca^2+^-dependent lysosomal fusion to the plasma membrane and the endosomal sorting complexes required for transport (ESCRT) complex-dependent membrane scission, have been extensively studied in multiple eukaryotic systems (1–4, 7–11). In contrast, the PMD-induced cell death response, pyroptosis, has been studied mostly in mammalian cells and the context of the immune response or cancer treatment (12, 13). Although these studies revealed the molecular mechanisms underlying the survival or death response after PMD in each system, a unified view of PMD response in eukaryotes remains largely elusive partly because of the lack of a universal PMD induction method. In particular, the PMD-inducing chemicals utilized in mammalian cells such as Streptolysin O or perforin cannot be easily adopted to cell types with a rigid cell wall including yeast.

Yeast serves as an excellent genetic tool to comprehensively identify genes required for fundamental cellular processes in eukaryotes. Previously, we demonstrated that budding yeast is equipped with a mechanism for repairing laser-induced PMD (14, 15). Although laser damage is a universal PMD method applicable for both budding yeast and higher eukaryotes, it cannot be easily employed in large-scale analysis.

Here, we developed a simple PMD induction method that can be utilized in budding yeast and human cultured cells. Using the assay, we performed a systematic yeast genetic screen and unexpectedly find that the genes required for PMD response and replicative lifespan regulations overlap significantly. Moreover, we find that PMD limits replicative lifespan in budding yeast and induces premature senescence in normal human fibroblasts. PMD-dependent premature senescence in normal human fibroblasts is mediated by the Ca^2+^-p53 axis, but not the best-studied senescence pathway, the ATM/ATR pathway. Our study proposes that PMD limits cellular lifespan in two different eukaryotic cell types. PMD-dependent senescence may explain the origin of senescent cells around the cutaneous wounds in vivo (16).

## Results

### Establishing a simple and universal PMD-inducing method

To reveal the conserved features of PMD response in eukaryotes, we needed a simple and reliable PMD-inducing method that can be 1) applicable for both budding yeast and human cells, and 2) utilized in large-scale analyses. A candidate chemical was sodium dodecyl sulfate (SDS) because previously we found that the wild-type budding yeast cells can grow on the YPD plate containing 0.02% SDS but the yeast mutants defective in PMD response fail to grow on it (14, 15). We tested whether SDS breaks cell wall and plasma membrane, which would lead to the penetration of a scarcely membrane-permeable fluorescent chemical, 4’,6-diamidino-2-phenylindole (DAPI). The DAPI goes into only 9.8±3.9% of wild-type budding yeast cells, consistent with the fact that the wild-type budding yeast can survive in the presence of SDS. However, in combination with EGTA, which prevents Ca^2+^-dependent membrane resealing (8), DAPI penetrated 74.5±4.9% (mean±SD) of cells (Fig. 1A and B). These results suggest that SDS breaks the plasma membrane, and the damage is immediately resealed by Ca^2+^-dependent mechanisms.

**Fig. 1.**
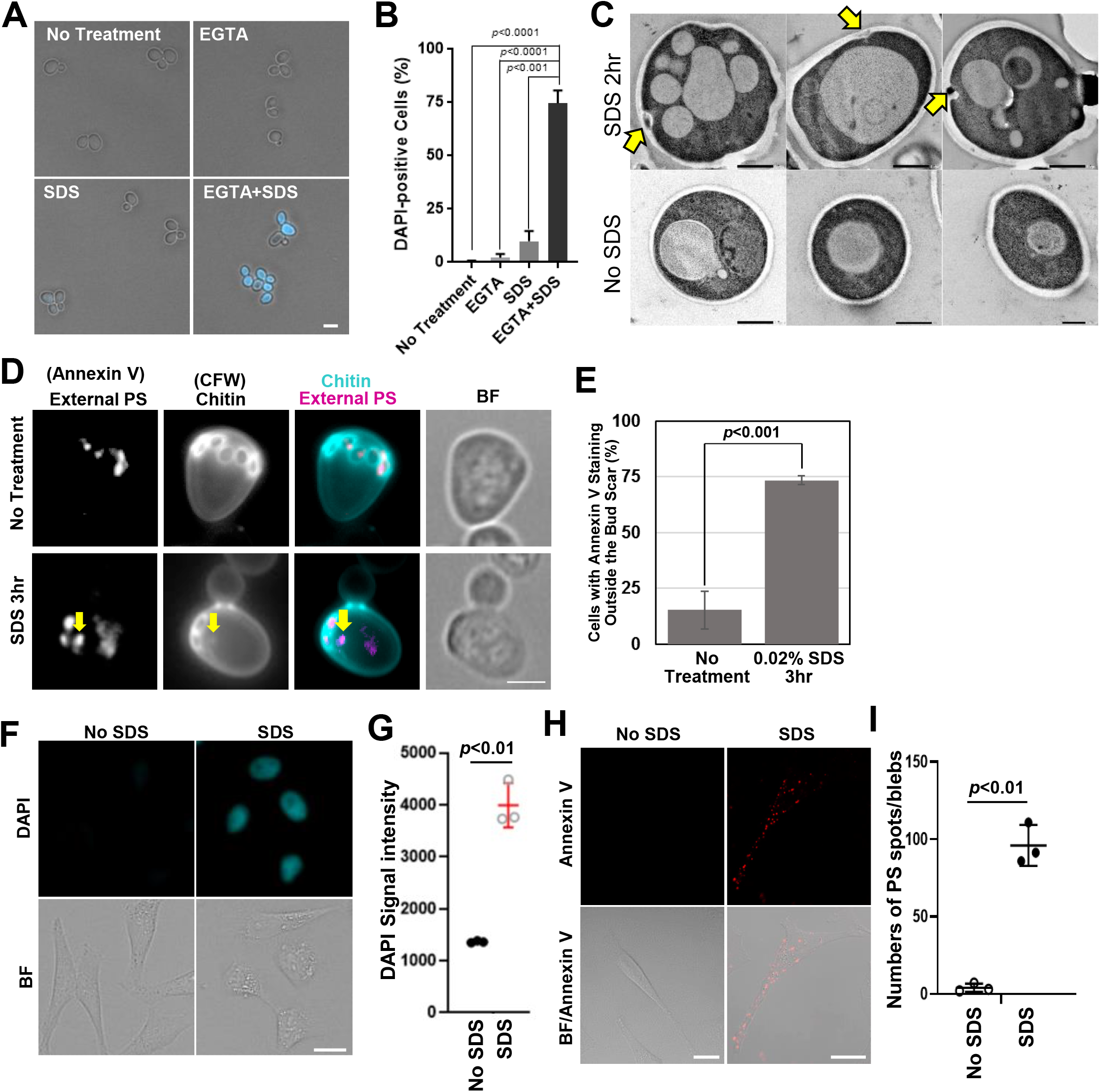
SDS induces PMD in yeast and human cells. **(A)** Wild type yeast cells were cultured in YPD and then incubated with 2 ng/ml DAPI-containing YPD with or without 20 mM EGTA and 0.02% SDS for 15 min. Scale bar, 5 μm. **(B)** Quantification of (A). Data are presented as the mean±SD of 3 independent experiments. N>300 cells/each experiment. *p*-value, by 2-tailed unpaired Student’s *t*-test. **(C)** Wild type yeast cells were cultured at 25°C and then incubated with 0.02% SDS for 2 hr. Cells were fixed and observed by the TEM. Yellow arrows, plasma membrane and cell wall ingression. Scale bar, 1 μm. **(D and E)** Wild type yeast cells were cultured in YPD and then switched to YPD containing 0.02% SDS for 3hr. Yeast cells were treated with zymolyase in 1.2 M sorbitol for 90 min and then stained with Annexin V-Alexa568 for 20 min. Yellow arrows, Annexin V and calcofluor white positive spots. % of cells with Annexin V staining outside of the bud scar was counted. Data are presented as the mean±SEM of 3 independent experiments. N>250 cells/each experiment. *p*-value, by 2-tailed unpaired Student’s *t*-test. **(F)** Representative images of DAPI penetration upon 0.008% SDS treatment for 24hrs. HeLa cells were cultured in DAPI-containing DMEM with or without 0.008% SDS. Scale bar, 20 μm. **(G)** Average DAPI intensity of three independent experiments as in (F). Untreated control and 0.008% SDS treatment were significantly different (*p*<0.01) using multiple Welch’s *t*-test with Benjamini, Krieger and Yekutieli correction. See also Fig. S2. **(H)** Representative images of the cells with Annexin V-positive spots upon 0.008% SDS treatment. HeLa cells were cultured in DMEM with or without 0.008% SDS for 1 hr. Scale bar, 20 μm. AnV-647: Annexin V, Alexa Fluor 647 conjugate, BF: Bright Field, Merge: Merged images of Annexin V and Bright Field. **(I)** Average numbers of Annexin V-positive spots of three independent experiments as in (H). See also Fig. S3A.

To investigate whether SDS also damages the cell wall, we observed chitin that staunches the laser-induced cell wall damage (14). We found local chitin spots in 84.9±5.5% of cells after SDS treatment (Fig. S1A, white arrows), analogous to the phenotype after laser damage (Fig. S1B, yellow arrows). Consistently, transmission electron microscopy images showed that SDS treatment induced local ingression of the cell wall and plasma membrane structure (Fig. 1C, yellow arrows). These results suggest that SDS breaks both cell wall and plasma membrane, and the damage is local, forming individual spots.

Next, phosphatidylserine (PS) externalization was examined because PMD leads to local PS externalization in human cells (17). Unexpectedly, PS was externalized at the bud scars, former cytokinesis sites marked by circular chitin staining in budding yeast, prior to the SDS treatment (Fig. 1D, upper panels). In addition to the signal at the bud scars, PS externalizing spots increased after SDS treatment, and the PS spots colocalized with chitin spots (Fig 1D, lower panels, and E). Moreover, two major repair proteins, Pkc1-GFP and Myo2-GFP, were recruited locally, but not globally, to the cortex after SDS treatment (Fig. S1C, yellow arrows). Altogether, these results demonstrate that SDS induces local plasma membrane and cell wall damage in budding yeast.

Next, we examined whether SDS damaged the plasma membrane of human cultured cells. We found that SDS treatment induced the influx of fluorescent dyes, DAPI and FM1-43, into HeLa cells (Fig. 1F and G, Fig. S2A-E). In addition, the PS-externalizing spots at the cell periphery increased after SDS treatment in HeLa cells (Fig. 1H and I, Fig. S3A and B), analogous to the treatment with other membrane-poring reagents (17). Live-cell imaging confirmed that PS-externalizing spots and blebs increased after SDS treatment in WI-38 cells (Fig. S3C). ESCRT-III (CHMP4A) signals were detected at the PS-externalizing spots/blebs (Fig S3D). These results indicate that SDS treatment induces PMD in human cells. Using this simple treatment, we designed a genome-wide screen to identify factors required for PMD response in budding yeast.

### Identification of PMD response genes in budding yeast

To identify genes essential for PMD responses, we performed a genome-wide screen using two yeast libraries: a non-essential gene deletion library (18), and a DAmP library in which mRNA levels of essential genes are decreased to 20–50% (19). These two libraries account for 96% of all open reading frames in budding yeast. We identified 48 mutants that were reproducibly sensitive to SDS (Fig. 2A, Fig S4A and B). The screening hits could be manually classified into 19 functional groups. The largest group in the hits was ESCRT with eight genes (Fig. 2B). GO enrichment analysis (http://geneontology.org) revealed that the cellular processes associated with ESCRT were highly enriched (Fig 2C and D, Table S1). Thus, ESCRT’s cellular function is essential for PMD response in budding yeast analogous to previously shown in higher eukaryotes (1–4, 7–11). The characterization of other screening hits is described in supplementary text (Supplementary text, Fig S5A-K and S6).

**Fig. 2.**
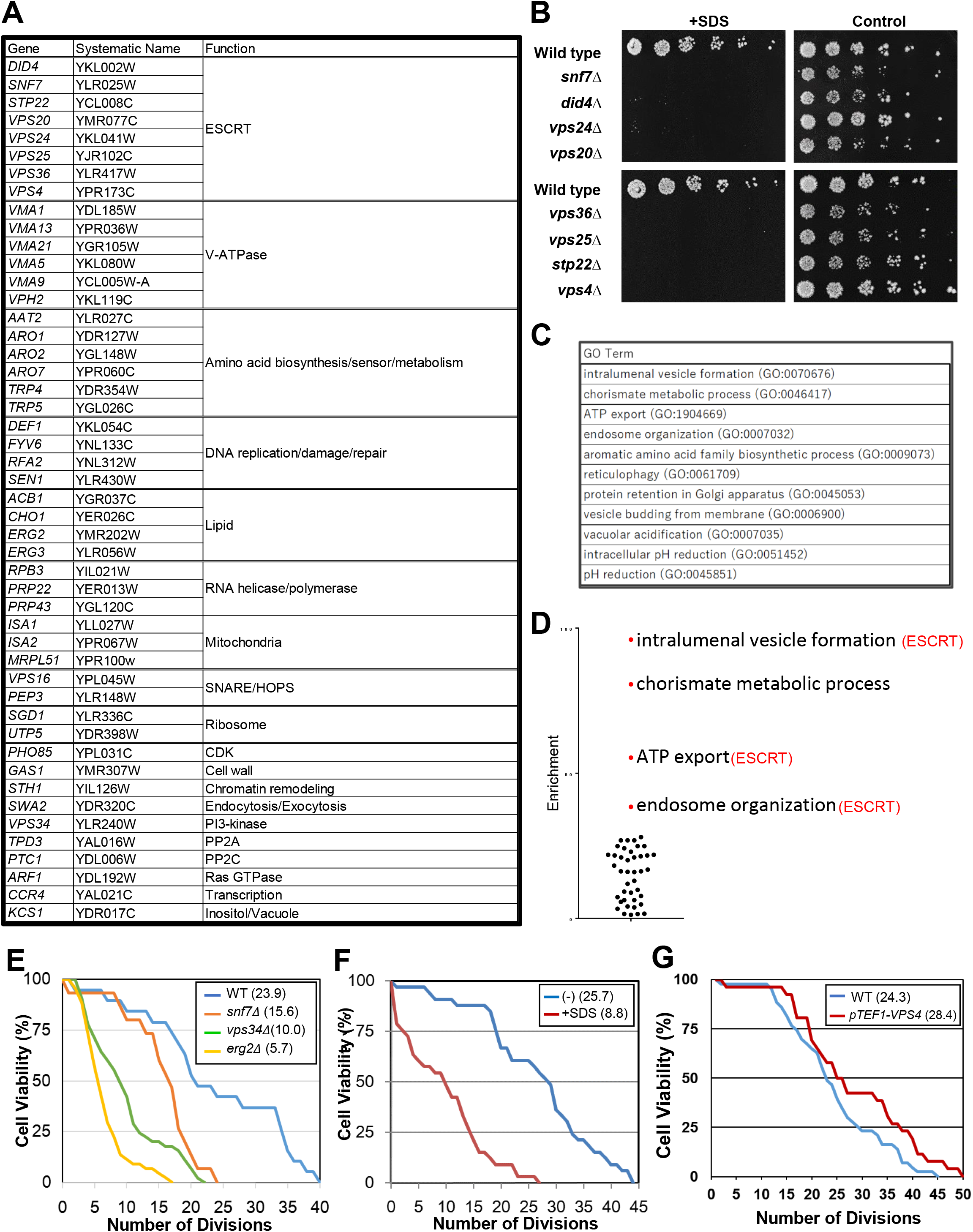
Identification of genes required for PMD response in budding yeast; PMD limits replicative lifespan in budding yeast. **(A)** The genes required for the growth in the presence of 0.02% SDS. **(B)** ESCRT mutants were spotted on YPD (4x dilution) with or without 0.02% SDS, incubated at 25°C for 3 days. **(C)** The enriched GO terms. See also Table S1. **(D)** Fold enrichment scores in the GO enrichment analysis were plotted. **(E)** Wild type, *snf7*Δ, *vps34*Δ, and *erg2*Δ were subjected to the replicative lifespan measurement. Mean values in parentheses. *p*-values: wild type and *snf7*Δ, *p*<0.01; wild type and *vps34*Δ, *p*<0.01; Wild type and *erg2*Δ, *p*<0.01 by 2-tailed unpaired Student’s *t*-test. **(F)** Wild type cells were subjected to the replicative lifespan measurement with or without 0.02% SDS. Mean values are shown in parentheses. *p*<0.01 by 2-tailed unpaired Student’s *t*-test. **(G)** Wild type or *VPS4*-overexpressing yeast cells were subjected to the replicative lifespan measurement. Mean values in parentheses. *p* value: 1-25 division, *p*=0.39; 26-50 division, *p*<0.05, by 2-tailed unpaired Student’s *t*-test.

### PMD limits the replicative lifespan of budding yeast

The gene sets required for the cellular/subcellular processes after PMD should be enriched in our hits. Therefore, to identify the reported phenotype enriched for our hits, we performed the model organism Phenotype Enrichment Analysis (*mod*PhEA) (20, http://evol.nhri.org.tw/phenome2). We found that the genes associated with the phenotype “replicative lifespan” were significantly enriched (Table S2 and S3). Motivated by this finding, we performed a replicative lifespan analysis (21) using our screening hits. All three mutants tested (*snf7Δ*, *vps34Δ*, and *erg2Δ*) showed no growth in the presence of SDS, which is consistent with our screening strategy (Fig. S4). *snf7Δ*, *vps34Δ*, and *erg2Δ* cells showed markedly shorter replicative lifespan, in the absence of SDS, compared with wild type cells (Fig. 2E), further suggesting a link between the PMD responses and replicative lifespan regulation. Next, we examined the replicative lifespan of wild type cells in the presence or absence of SDS. We found that SDS significantly shortened the replicative lifespan (Fig. 2F). Consistent with these results, overexpression of the ESCRT-activator AAA-ATPase *VPS4* extended the replicative lifespan (Fig. 2G). These results suggest that PMD limits the replicative lifespan of budding yeast, and ESCRT’s function is critical for the regulation of replicative lifespan.

### Transient PMD induces premature senescence in normal human fibroblasts

To test the possibility that PMD induces premature senescence in human cells, we performed a long-term culture of normal human fibroblasts (BJ and WI-38) in the presence or absence of PMD. Indeed, cell proliferation was inhibited in an SDS concentration-dependent manner (Fig. 3A and B).

**Fig. 3.**
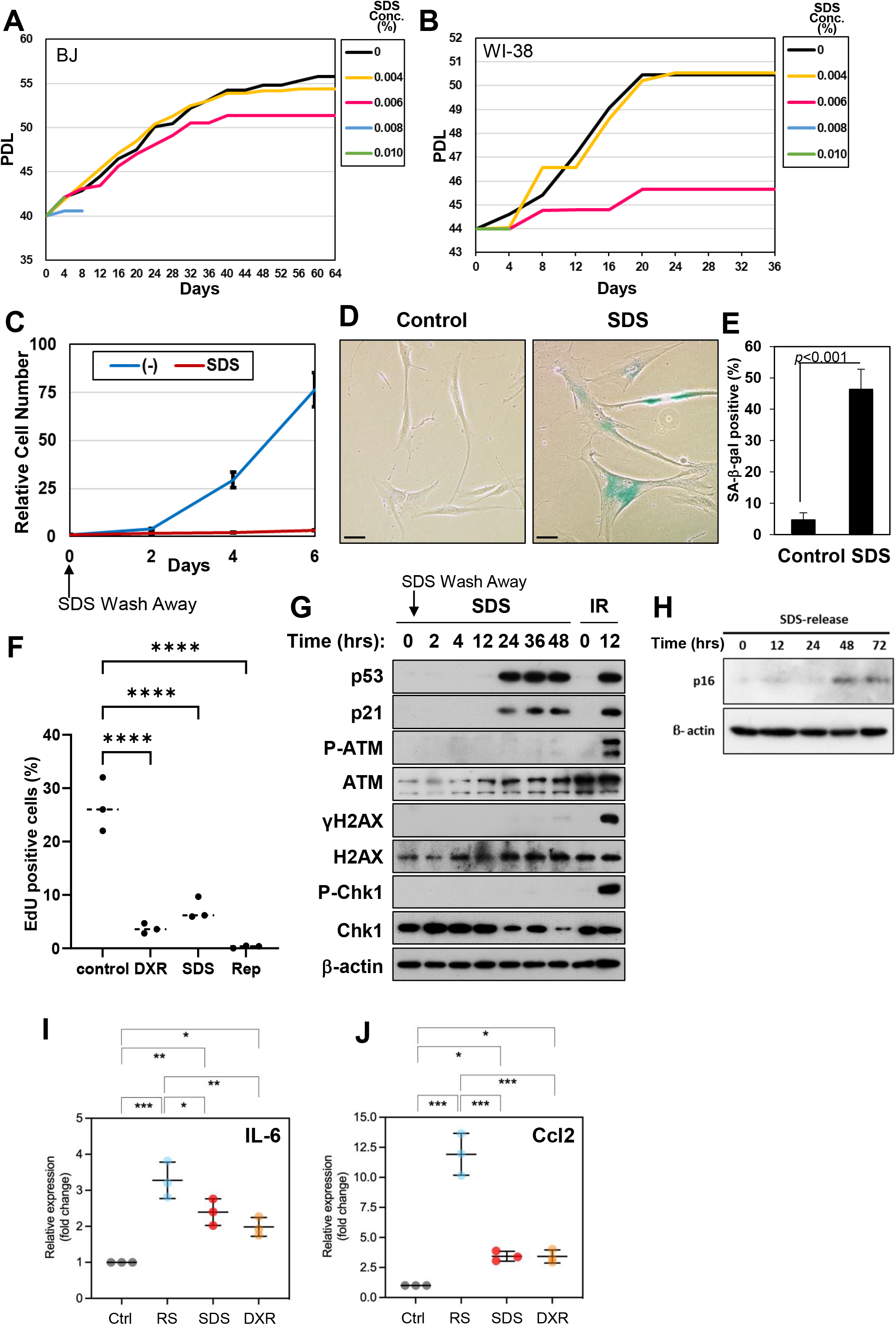
Transient PMD induces premature senescence in normal human fibroblasts. **(A and B)** Human normal fibroblasts (A: BJ, B: WI-38) were cultured with or without SDS (0.006-0.01%; indicated on the right). **(C)** HCA-2 cells were incubated with medium with (SDS) or without (-) 0.008% SDS for 24 hr, washed, and released into fresh medium. Data are presented as the mean±SD of 3 biological replicates. **(D and E)** HCA-2 cells were treated with 0.01% SDS for 24 hr, washed, and released into fresh medium. SA-β-gal-positive cells were detected using the cells 6 days after SDS wash away. Scale bar, 50 μm. Graphs show quantification of SA-β-gal-positive cells (n>100). Data are presented as the mean±SD of >3 independent experiments. *p*-value, by 2-tailed unpaired Student’s *t*-test. **(F)** WI-38 young cells (control) and senescent cells were labelled with 10 μM 10 μM EdU for 24 hrs. EdU-coupled Alexa Fluor 647 signals and Hoechst 33342 signals were obtained and the ratio of EdU-incorporated cells were calculated. The data are presented as means SD of 3 independent experiments. >200 cells per each experiment. *p*-value: **** < 0.001 by One-way ANNOVA, Dunnett.**(G and H)** Cell lysates of HCA-2 cells treated with 0.01% SDS for 24 hr or 10 Gy IR (Irradiation, control) at the indicated times after SDS wash away or IR treatment, respectively, were subjected to immunoblotting using the antibodies indicated. **(I and J)** The expression of the senescence-associated secretory phenotype (SASP) marker genes (H: IL-6, I: Ccl2) in senescent WI38 cells were analyzed by qPCR. WI-38 cells were treated with 0.0095% SDS for 24 hr (SDS) or 250 ng/ml doxorubicin for 24 hr (DXR). RNA was isolated from these cells after 5 days. RNA was also isolated from Replicative senescent cells (RS) and untreated WI-38 cells (Ctrl). Gene expression was normalized to that of Actb and calculated as fold changes. Error bars, SD. * p < 0.05, **p<0.01, ***p<0.001, (n=3).

To minimize side effects, we transiently treated normal human fibroblasts with SDS. HCA-2 and WI38 cells were incubated with SDS-containing media for 24 hrs, washed with medium, and then cultured in fresh medium. As a result, cell proliferation was inhibited after the treatment, and the proportion of senescence-associated ß-galactosidase (SA-ß-gal)-positive cells was increased after six days (Fig. 3C-E, Fig. S7A and B). EdU incorporation was attenuated in the cells 16 days after PMD treatment, indicating that DNA replication halted (Fig. 3F). Consistently, the protein levels of p53 (Fig. 3G and Fig S7C), an essential senescence regulator in mammalian cells (22, 23), and its target, p21, increased in cells treated with SDS as early as 24 hrs after SDS wash away (Fig. 3G). The levels of p16*INK4A* also increased 48 hrs after the SDS wash away (Fig. 3H and Fig S7C). The mRNA levels of SASP factors, *IL-6* and *CCL2*, were upregulated in SDS-treated cells analogous to replicative senescent and DNA damage-treated cells (Fig. 3I and J). Altogether, PMD triggers premature senescence in normal human fibroblasts.

### The plasma membrane damage-dependent senescence is suppressed by transient overexpression of ESCRT-III, CHMP4B

If SDS induces cellular senescence via plasma membrane damage, it should be suppressed by upregulation of plasma membrane repair. To test this assumption, we examined whether transient overexpression of ESCRT-III, CHMP4B, attenuates PMD-induced senescence. We transfected the plasmid harboring GFP-CHMP4B into WI-38 cells and the cells were treated with SDS as described above. We found that overexpression of CHMP4B bypassed SDS-induced proliferation arrest and decreased the proportion of SA-β-gal-positive cells (Fig. 4A-C). Consistent with these results, SDS-induced upregulation of p53, p21, and p16 protein levels was attenuated in the CHMP4B-expressing cells (Fig. 4D and E). The SASP factors upregulation was suppressed as well (Fig. 4F). Altogether, these results are consistent with our understanding that SDS induces premature senescence via plasma membrane damage and upregulation of plasma membrane repair suppresses it.

**Fig. 4.**
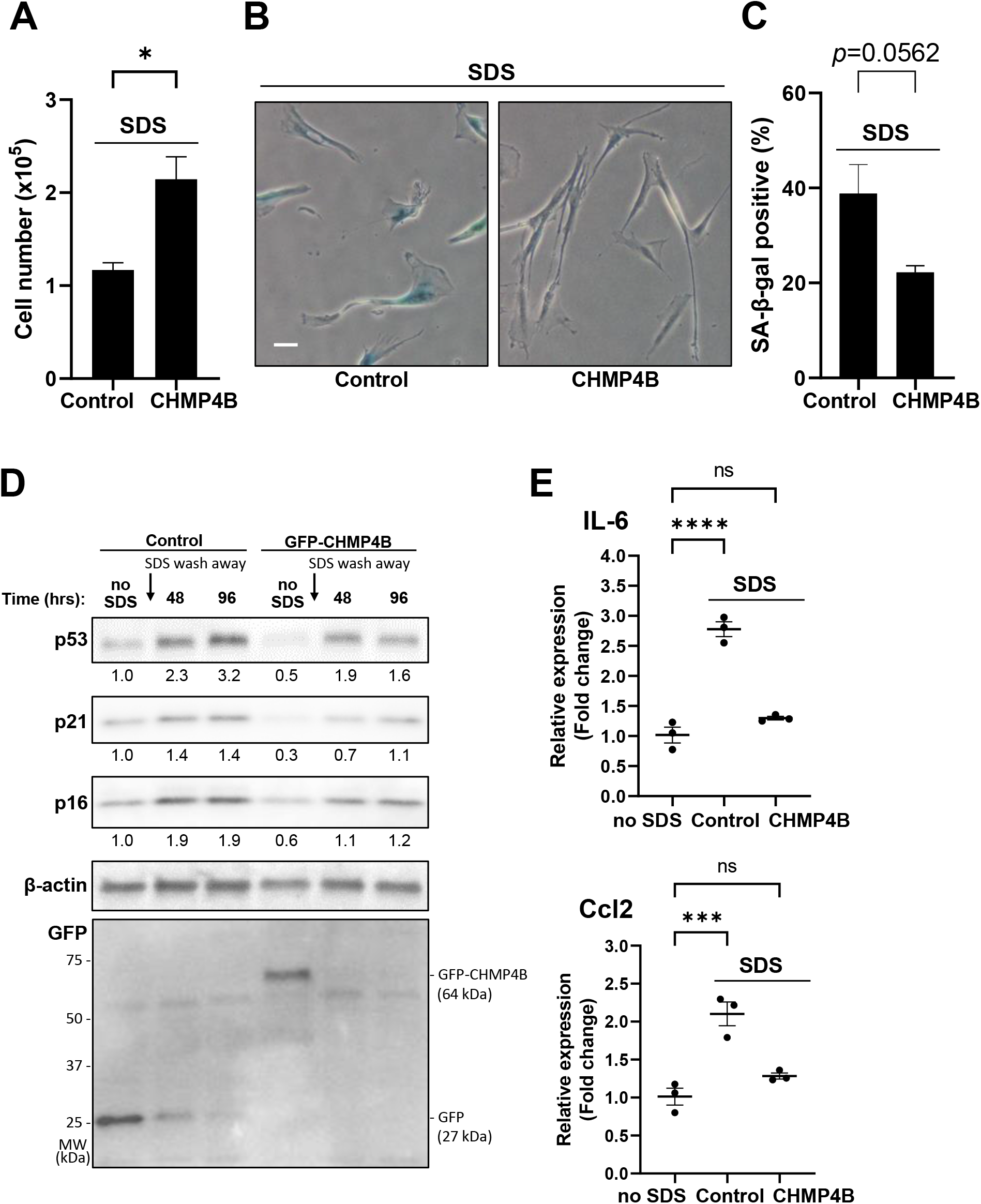
Transient overexpression of ESCRT-iii (CHMP4B) suppressed PMD-induced premature senescence. **(A-E)** A control plasmid (GFP) or a plasmid encoding ESCRT-iii (GFP-CHMP4B) were overexpressed in WI-38 cells. Cells were treated with 0.007% SDS for 24 hr, washed, and released into fresh medium. **(A)** Cell number in 6 cm diameter dish was counted 10 days after SDS wash away. *p*-value: * < 0.05 by student’s *t*-test. **(B and C)** SA-β-gal-positive cells were detected seven days after SDS wash away. Scale bar, 50 μm. Graphs show quantification of SA-β-gal-positive cells (n>100). *p*-value, by Student’s *t*-test. **(D)** Western blot was performed as in Fig. 3(G). GFP (control) and GFP-CHMP4B expression was confirmed by anti-GFP blot. Relative signal intensities are quantified and indicated under the blots. **(E)** qPCR was performed as in Fig. 3(I). p-value: *** < 0.005, **** < 0.001 by One-way ANNOVA, Dunnett.

### PMD induces cellular senescence via p53 in normal human fibroblasts

The best-characterized cellular senescence mechanism is ATM/ATR-dependent upregulation of p53-p21, which is activated after many senescence-inducing stimuli, including telomere shortening, DNA damage, and oncogene activation (24, 25). We tested whether this pathway is activated after PMD. Importantly, SDS increased the protein levels of p53 and p21, but not phospho-ATM, γH2AX, or phospho-Chk1 (Fig. 3G). Consistently, immunostaining with the γH2AX antibody was negative in the PMD cells at 24 hrs after the SDS wash away (Fig. 5A). These results suggest that the ATM/ATR pathway is dispensable for PMD-dependent upregulation of p53-p21.

**Fig. 5.**
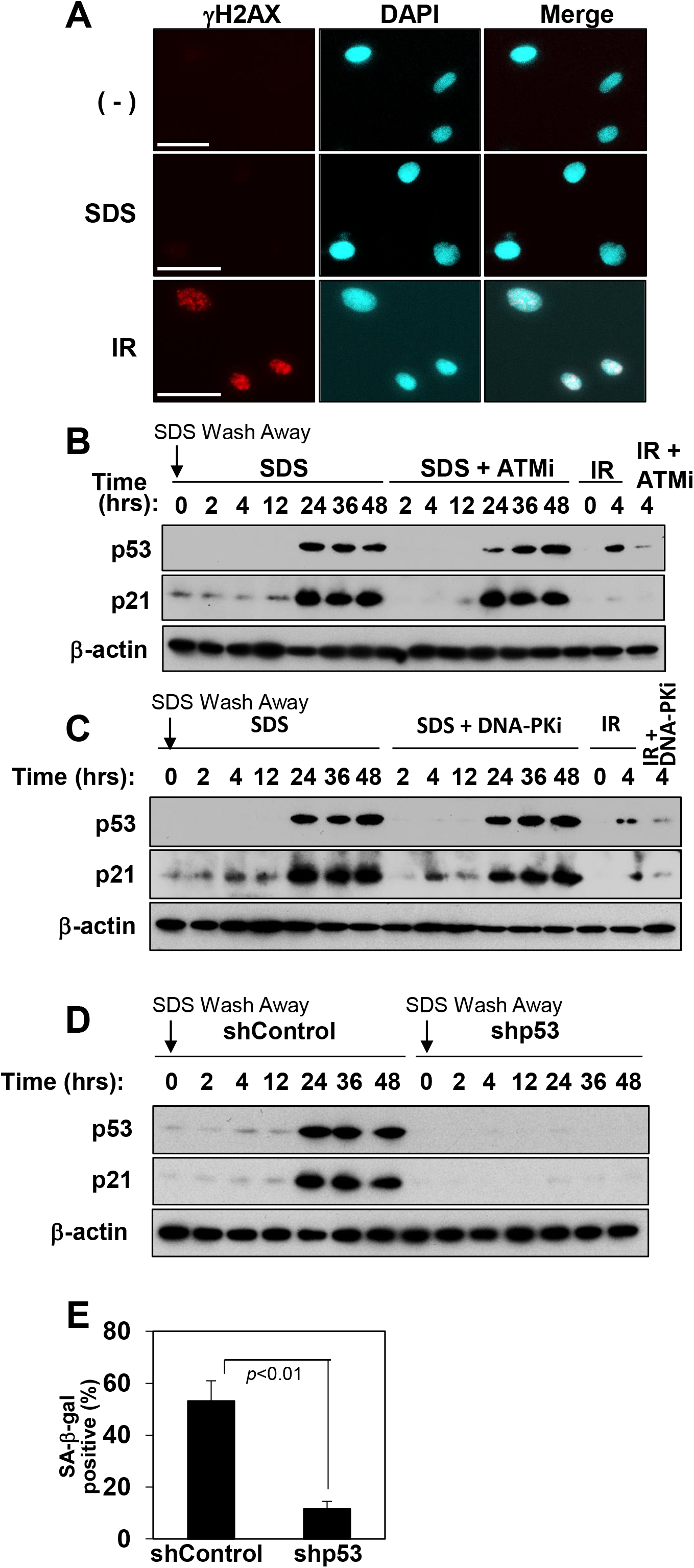
PMD-dependent premature senescence requires p53. **(A)** HCA-2 cells at 24 hr after SDS wash away (SDS), cells at 4 hr after 10 Gy Irradiation treatment (IR), and untreated cells (-) were stained with γH2AX (red) and with DAPI (blue). Scale bar, 50 μm. **(B and C)** HCA-2 cells were treated as in Fig. 3F with or without (B) ATM inhibitor KU-55944 (10 μM) or (C) DNA-PK inhibitor (10 μM). **(D)** Cells expressing Dox-inducible control shRNA (shControl) or p53 shRNA (shp53) with doxycycline (1 μg/ml) were treated with 0.01% SDS for 24 hr. After SDS wash away, cells lysates prepared at the indicated times were subjected to immunoblotting using the antibodies indicated. **(E)** SA-β-gal-positive cells were counted on 6 days after SDS wash away. *p*-value, by 2-tailed unpaired Student’s *t*-test.

To test this possibility further, HCA-2 cells were treated with the inhibitors of ATM and DNA-PKs (Fig. 5B and C). Both inhibitors did not alter SDS-dependent upregulation of p53 and p21. In contrast, p53-knockdown abolished p21 induction (Fig. 5D) and significantly decreased the proportion of SA-β-gal-positive cells at 6 days after SDS wash away (Fig. 5E). Complementarily, doxorubicin did not induce detectable plasma membrane rupture in the DAPI penetration assay (Fig. S8A and B). Altogether, we conclude that PMD induces cellular senescence via p53 but not ATM/ATR.

### Cytosolic Ca^2+^ levels increase after PMD

Ca^2+^ influx is one of the earliest responses after PMD in all cell types (1,2), and Ca^2+^ signaling is involved in cellular senescence (26). Therefore, we tested whether cytosolic Ca^2+^ increases after SDS treatment. We found that Cytosolic Ca^2+^ levels increased as early as one min after the SDS addition (Fig. 6A-C). Enforced Ca^2+^ influx by 75 mM KCl for 24 hrs was sufficient for increasing SA-β-gal-positive cells as well as p53, p21, and p16 protein levels (Fig. 6D-F). These results support our idea that Ca^2+^ influx plays an important role in p53 induction during PMDS (Fig. 6G).

**Fig. 6.**
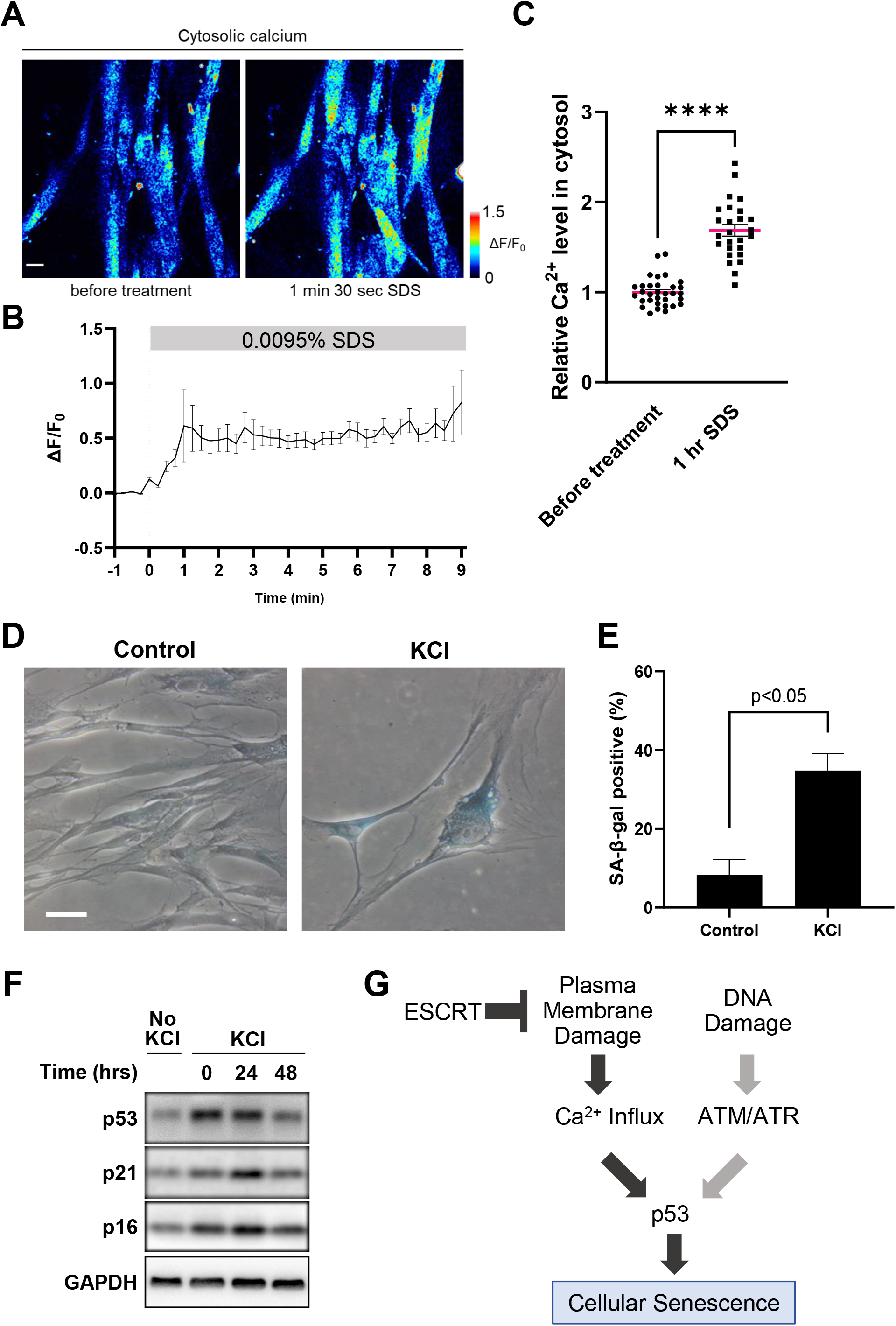
KCl-dependent Ca^2+^ influx is sufficient to induce premature senescence. **(A)** Representative images of cytosolic Ca^2+^ level (Cal Red R525/650) in WI-38 cells before or 1.5 min after 0.0095% SDS addition. Fluorescent intensity was coded by rainbow pseudo-color. Scale bar, 20 μm. **(B)** Representative traces of Cal Red R525/650 signals in WI-38 cells with 0.0095% SDS treatment. N=8. **(C)** Cytosolic Ca^2+^ level in WI-38 cells before or 1 hour after 0.0095% SDS treatment. N>25. Error bars, SEM. **(D and E)** WI-38 cells were treated with 75mM KCl for 24 hr, washed, and released into fresh medium. SA-β-gal-positive cells were detected using the cells seven days after KCl wash away. Scale bar, 50 μm. Graphs show quantification of SA-β-gal-positive cells (n>100). Data are presented as the mean±SEM of 3 independent experiments. p-value, by 2-tailed unpaired Student’s t-test. **(F)** WI-38 cells were treated with 75mM KCl for 24 hr, washed, and released into fresh medium. Samples were collected at indicated time points. Western blot was performed as in Fig. 3(G). **(G)** Summary of the PMD-dependent cellular senescence mechanism.

### PMD-dependent senescent cells (PMDS) and replicative senescent cells have PS-externalizing spots/blebs, but DNA damage-dependent senescent cells do not

Since PS-externalizing spots/blebs were observed after PMD (Fig. 1H and I, Fig. S3A-D), we determined whether such spots/blebs exist in senescent cells. As expected, Annexin V-positive PS-externalizing spots/blebs were detected in PMDS (Fig. 7A, SDS). Intriguingly, the replicative senescent cells, but not IR-treated senescent cells, had the Annexin V spots/blebs (Fig. 7A, RS and IR). Analogous to the PMD cells, ESCRT-III CHMP4A signals were detected at 21.5±5.0% (mean±S.E.M.) of PS-externalizing spots/blebs in replicative senescent cells (Fig. 7B). Some PS spots colocalize with projection-like structures in the bright field image (Fig. 7A, white arrows); therefore, we performed a focused ion beam scanning electron microscope (FIB-SEM) analysis. We found that untreated young normal human fibroblasts (HCA-2) had smooth plasma membranes (Fig. 7C, upper). In contrast, PMDS had projections (Fig. 7C, lower). The heights of the projections were diverse (280 nm to 2.5 μm). The widths of the projections were relatively uniform (100–130 nm). Altogether, these results demonstrate that PS-externalizing projections are the common feature of PMDS and replicative senescent cells, but not of DNA damage-dependent senescent cells.

**Fig. 7.**
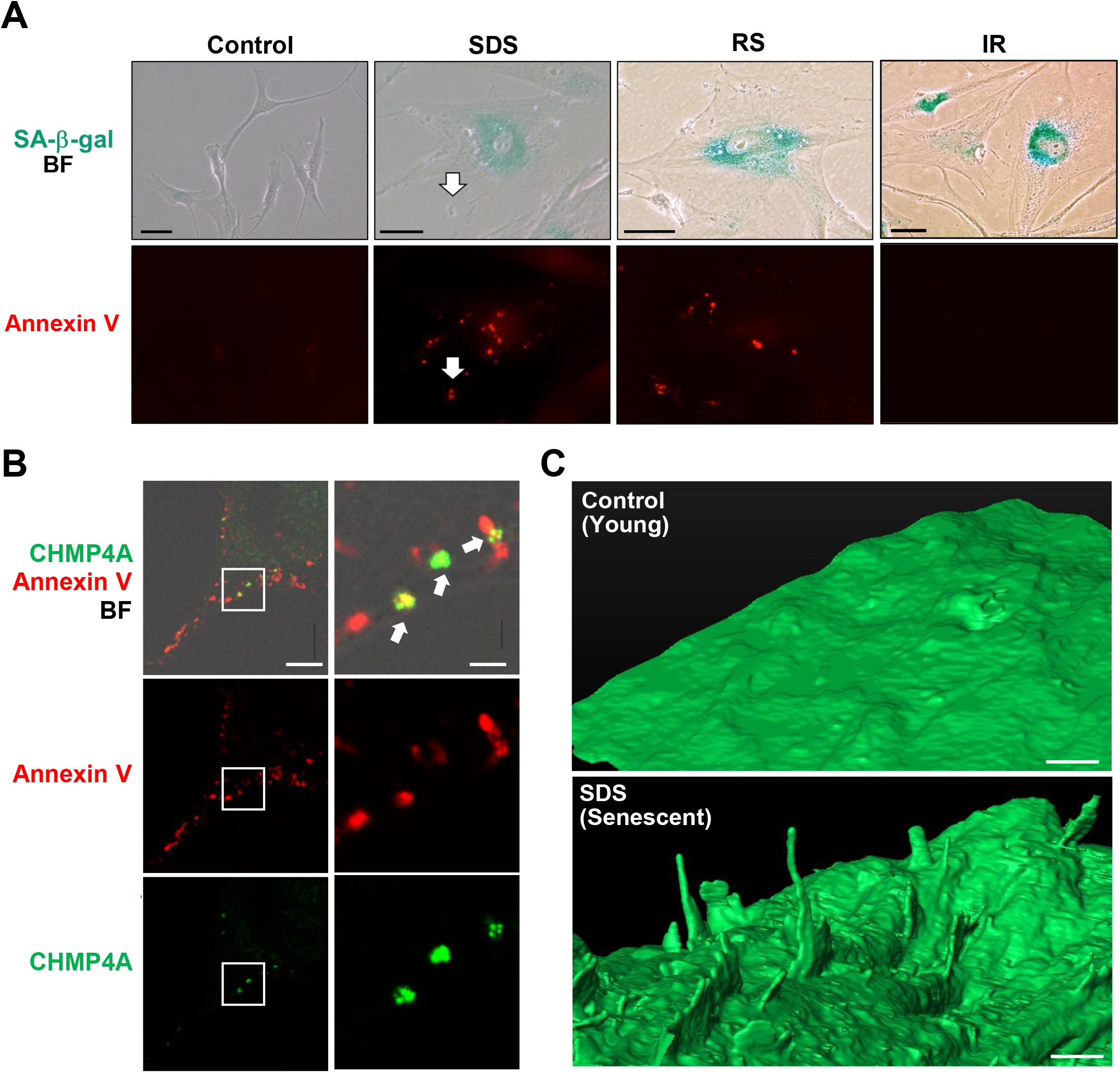
Local PS externalization is associated with PMD-dependent and replicative senescence. **(A)** SA-β-gal-positive (green) and Annexin V-Alexa 488-positive (red) HCA-2 cells were identified using the cells 6 days after SDS wash away (SDS), replicative senescence (RS), or the cells 6 days after irradiation (IR). Scale bar, 50 μm. White arrows, an Annexin V-positive projection. **(B)** Replicative senescent WI-38 cells were incubated with Annexin V-Alexa Fluor 647 conjugate, fixed, and stained with CHMP4A antibody. The white rectangle region in left panels is enlarged in right panels. Green: CHMP4A, Red: Annexin V, Alexa Fluor 647 conjugate, BF: Bright Field. White arrows indicate Annexin V and CHMP4A colocalization. Scale bars, left; 10 μm, right; 2 μm. **(C)** HCA-2 cells were treated with 0.01% SDS for 24 hr, washed, and released into fresh medium. The cells were fixed after 6 days. SDS-treated (SDS, Senescent) and untreated (Control, Young) cells are analyzed by FIB-SEM. Scale bar, 1 μm.

## Discussion

Here, we report that PMD limits replicative lifespan in budding yeast and induces stress-dependent premature senescence in human normal fibroblasts. We have developed a simple and universal PMD-inducing method and designed a systematic genome-wide screen using budding yeast. Based on the screen, we find that PMD limits replicative lifespan in budding yeast, but the overexpression of *VPS4* extends it. In normal human fibroblasts, PMD induces premature senescence mediated by p53. Our work demonstrates that cellular senescence is a universal but underappreciated cellular consequence of PMD in eukaryotes.

### ESCRT activation associated with replicative lifespan extension in budding yeast

ESCRT is involved in various cellular processes, including cytokinesis, plasma membrane repair, nuclear membrane repair, multivesicular body formation, vacuolar membrane repair, autophagosome formation, and microautophagy of the endoplasmic reticulum (11, 27, 28). Here, we find ESCRT in our screen for PMD response factors. Moreover, we show that ESCRT is important for budding yeast replicative lifespan regulations based on two lines of evidence: 1) a yeast mutant lacking an ESCRT-III component (*snf7Δ*) showed a short replicative lifespan (Fig. 2E), and 2) *VPS4*-overexpression extended the replicative lifespan of budding yeast (Fig. 2G).

ESCRT can contribute to the replicative lifespan by at least two scenarios that are not mutually exclusive. First, ESCRT-dependent plasma membrane repair may prevent cell lysis and extend the replicative lifespan through the ESCRT’s function in plasma membrane repair as established in human cells (11, 28) and recently suggested in yeast (28). Second, ESCRT-dependent vacuolar membrane repair may extend the replicative lifespan of budding yeast. This explanation also makes sense because vacuolar acidification defects limit the replicative lifespan in budding yeast (30, 31). ESCRT is involved in nuclear envelope repair (11). However, we found that SDS treatment did not induce marked nuclear envelope deformation (Fig. 1F and Fig. S2A); ESCRT did not accumulate at the nuclear membrane after SDS treatment (Fig. S3D, green). These results suggest that the SDS-dependent cellular senescence is not via the nuclear envelope damage. Our study demonstrates that the PMD response shares many regulators, including ESCRT, with replicative lifespan regulation, and that incomplete PMD response explains a mechanism underlying DNA damage-independent short replicative lifespan in budding yeast.

### Budding yeast and human cells share part of stress-dependent cell cycle arrest mechanisms

Budding yeast has been utilized as a model of replicative lifespan regulations in eukaryotic cells. In telomere biology, at least three features are common between budding yeast and human cells: 1) The telomerase deficiency induces senescence in budding yeast (32), 2) Telomere shortening causes senescence via permanent activation of DNA damage checkpoints both in budding yeast and human cells (33), and 3) Telomere shortening-dependent DNA damage checkpoints can be bypassed due to adaptation both in human cells and budding yeast (34, 35). Thus, yeast can serve as a model to study basic senescence mechanisms associated with cell cycle regulations. However, there are limitations. For example, yeast does not have the key senescence regulators such as p53 and SASPs. Thus, yeast can serve as a model to study senescence-triggering cell cycle checkpoints but not all aspects of cellular senescence mechanisms.

### Molecular mechanisms underlying PMD-dependent replicative lifespan shortening

We previously reported that PMD induces cyclin-dependent kinase (CDK) inhibitor Sic1 stabilization in budding yeast (15). Analogous to budding yeast, here we show that PMD promotes upregulation of two CDK inhibitors, p21 and p16^*INK4A*^, in normal human fibroblasts (Fig. 3G, H, and Fig. S7).

The next question is how PMD triggers the CDK inhibitor upregulation. We demonstrate that p53 was essential for the upregulation of p21 in normal human fibroblasts (Fig. 5D), but the ATM/ATR pathway was dispensable (Fig. 5B and C). Other group demonstrated that high cytosolic Ca^2+^ levels induce mitochondrial impairment, leading to upregulation of p53/p21 via cAMP-responsive element-binding protein (CREB) (36). Consistently, here we show that the cytosolic Ca^2+^ increases after the plasma membrane damage (1–3; Fig. 6A-C); enforced Ca^2+^ influx by KCl was sufficient for inducing various senescence features (Fig. 6D-F). Therefore, Ca^2+^ influx after PMD may mediate the upregulation of the p53-p21 axis in normal human fibroblasts. The absence of p53 in budding yeast suggests that the molecular details are not identical in budding yeast and human cells. Nonetheless, the upregulation of CDK inhibitors is a common mechanism promoting the PMD-dependent cell cycle arrest in both systems.

### Local PS externalization as a marker of specific senescent cell subtypes

Here, we observed PS-externalizing spots/blebs in PMD-dependent and replicative senescent cells, but they were absent from DNA damage-dependent senescent cells (Fig. 7A). Although PS externalization after PMD was reported (17), the origin of the PS-externalizing spots/blebs in the replicative senescent cells remains unknown. Plasma membrane repair and cytokinesis share the membrane resealing machinery, including ESCRT (3,11); therefore, repeated cytokinesis could induce the accumulation of the PS-externalizing spots/blebs during replicative senescence. This idea is consistent with our finding that PS was externalized at the bud scars, former cytokinesis sites, in budding yeast (Fig. 1D). Alternatively, the lipid scramblases (37) could be activated in replicative senescent cells. In either case, our results demonstrate that the accumulation of PS-externalizing spots/blebs is the feature of PMDS and replicative senescent cells.

Overall, our results highlight an underappreciated subtype of senescent cells. Recent studies demonstrated that senolytic drugs clearing senescent cells ameliorate age-associated disorders (38, 39). Our study, therefore, may serve as a basis for further studies aimed at developing new therapeutic strategies for PMD-associated diseases, including muscular dystrophy and Scott syndrome, as well as organismal aging.

## Materials and Methods

### Media, strains, and genetic manipulations

Standard procedures were employed for DNA manipulations, as well as for *E. coli* and *S. cerevisiae* handling. Yeast extract peptone dextrose (YPD) was used in most of experiments. SD was used for live-cell imaging. Yeast culture was performed at 25°C unless otherwise indicated. The *S. cerevisiae* strains used in this study are listed in Table S3.

### DAPI-penetration assay

A budding yeast overnight culture in YPD was refreshed and incubated for an additional 2-6 hours until the OD_600_ reached 0.1-0.3. For Fig. 1A and B, the media was switched to YPD+20 mM EGTA, YPD+0.02% SDS, YPD+20 mM EGTA+0.02% SDS, and control YPD, for 15 min. All of these media contained 2 ng/ml DAPI. DAPI-positive cells were counted under Celldiscoverer 7 (Zeiss).

### Fluorescence imaging and image analysis in yeast

The laser damage assay was performed as described previously (14) with modifications. A yeast culture grown overnight was refreshed and incubated for an additional 2-6 hours until the OD_600_ reached 0.1-0.3. Cells were then spotted onto an agarose bed (SD medium+1.2% agarose) on glass slides. LSM 780 NLO confocal laser scanning microscopy (Zeiss) was used to induce the damage and to monitor the fluorescent and bright-field images. Signal quantification was performed using FIJI software (40).

### Electron microscopy of yeast

Yeast cells sandwiched between copper grid were subjected to quick freezing using an isopentane/propane mixture of cold chemicals. The copper grid was diverged under liquid nitrogen gas atmosphere, soaked in 2% osmium/acetone solution, and then stored at −80°C for 2 days. The samples were placed at −20°C for 3 hours, at 4°C for 1 hour, and then at room temperature. The samples were washed with acetone, and subsequently with propylene oxide, and then embedded in epoxy resin (Quetol-651). Observation was performed under JEM-1011J (JEOL) or JEM-1400Plus (JEOL).

### Yeast genome-wide screen

SDS sensitivity was determined using freshly prepared YPD plates containing 0.02% SDS. Cells grown on YPD plates for 3 days were pin-plated onto the SDS-containing plates and incubated at 30°C for 3 days. The mutants identified in the first screening (249 mutants) were then freshly grown from −80°C stock. Single colonies were isolated, streaked on YPD and YPD 0.02% plates, and incubated at 30°C for 3-5 days. The incubation time was optimized for each strain. We obtained 109 mutants from the library, which were lethal on YPD+0.02% SDS plates. Subsequently, we independently constructed 119 mutants using our wild type (BY23849) as a parental strain. At least three independent colonies were constructed per each mutant. Single colonies were isolated, streaked on YPD and YPD 0.02% plates, and incubated at 25°C for 3-5 days (optimized for each strain). In the end, we obtained 48 mutants as confirmed hits, which were reproducibly lethal specifically on YPD+0.02% SDS plates.

### Replicative lifespan analysis in yeast

Replicative lifespan analysis was performed as described (21). Briefly, newly formed daughter cells were separated after every cell division by a glass needle for tetrad analysis under the microscope. During the night, the plates were incubated at 12°C.

### Co-staining of Annexin V-Alexa 568 and calcofluor white in yeast

Externalized PS detection in yeast was performed as described (41) with some modifications. Yeast cells were cultured in YPD for overnight, diluted to 1/10 in fresh YPD, and then incubated for additional 2-3 hours. Cells were collected and washed once with Sorbitol buffer (35 mM potassium phosphate, pH 6.8, 0.5 mM MgCl_2_, 1.2 M sorbitol). Cells were incubated in 98 μl Sorbitol buffer + 2 μl of 2.5 mg/ml Zymolyase 100T (SEIKAGAKU CORPORATION) for 90 min (wild type) or 60 min (*pTEF1-VPS4*) at room temperature with gentle shaking. Cells were then washed once with Sorbitol buffer, resuspended in 19 μl Annexin V binding buffer + 1 μl Annexin-V-Alexa 568, and incubated for 20 min at room temperature with gentle shaking. Supernatant was removed and cells were suspended in 9.8 μl Annexin V binding buffer + 0.2 μl Calcofluor White Stain (Sigma-Aldrich, Fluka). Images were acquired using Axio Observer z1 (Zeiss).

### Cell culture methods

Cell culture methods, senescence induction by irradiation and SA-β-gal staining were performed as described (22, 42). Immunoblotting was performed following a standard protocol. Antibodies used in this study were listed in Table S2. Normal human fibroblasts (HCA-2, WI-38, or BJ) were used. HCA-2 cells are neonatal foreskin cells (42). ATM inhibitor (KU-55933, Sigma-Aldrich) and DNA-PK inhibitor (NU7026, Sigma-Aldrich) were added to the medium at a final concentration of 10 μM. For Fig. 3A and B, cells were grown in the medium containing 0.006-0.01% SDS or no SDS (control), and passage was performed every 3-4 days. After every passage, the cells were grown in the medium without SDS for 16 hrs, and then switched to the SDS-containing medium. For SDS-dependent senescence induction, cells were treated with the medium containing SDS (0.0085-0.01%, the concentration was optimized for each experimental condition) for 24 hours, washed twice with medium and covered with the fresh medium, and then incubated for six additional days. During the six days, medium was changed every two days. For KCl treatment (Fig. S8A), WI-38 cells were treated with 75mM KCl for 24 hr, washed, and released into fresh medium. Annexin V-Alexa Fluor 488 (Thermo Fisher Scientific) staining was performed following the manufacturer’s instructions. Images were acquired using a BZ-9000 all-in-one fluorescence microscope (Keyence) or LSM-780 Confocal Microscopy (Zeiss). For time-lapse imaging (Fig. S3C), WI-38 cells were incubated with the medium containing 0.008% SDS and 1:5,000 diluted pSIVA-IANBD (Novus Biological). Images were acquired by Celldiscoverer 7 (Zeiss).

### CHMP4B expression in WI-38 cells

A control plasmid (GFP) or GFP-CHMP4B were overexpressed in WI-38 cells by 4D-Nucleofector (Lonza; SE cell line solution; program EO-114) according to the manufacturer’s instructions. One million cells were nucleofected with 0.5 μg DNA of either GFP or GFP-CHMP4B.

### Co-staining of AnnexinA5–Alexa647 and CHMP4A in human cells

HeLa cells were plated on a glass bottom dish (greiner bio-one) coated with 0.1% gelatine one day before the experiment. HeLa cells were treated with 0.008% SDS in growth medium (9% FBS DMEM) for 1h at 37°C. Replicative senescent WI-38 cells were plated on 0.1% gelatine coated coverslips (Mastunami) submerged in 6 well cell culture dish (VIOLAMO). Cells were washed twice with growth medium and externalized PS was stained with AnnexinA5–AlexaFluor-647conjugate (Invitrogen) diluted to 0.002% (v/v) in growth medium for 15 min at room temperature. Cells were washed twice with growth medium and then fixed with 4% paraformaldehyde (Electron Microscopy Sciences) for 15 min at 37°C. Cells were permeabilized and blocked in 1xPBS+ containing 0.1% saponin (Sigma) and 5% BSA (Sigma-Aldrich) for 40 min at room temperature. Subsequently cells were incubated overnight with primary antibody (Anti-CHMP4A antibody: Sigma-Aldrich HPA068473) in 1% BSA in 1xPBS+ at 4°C. Cells were washed four times with PBS+0.5% FBS and incubated with goat anti-rabbit IgG secondary AlexaFluor 488 conjugated antibody (Invitrogen A32731) for 1 h at room temperature. Cells were washed four times with PBS+ 0.5% FBS. For coverslip samples, coverslips were mounted on slide glass using Antifade Glass mounting medium (Invitrogen). Images were taken with SP8 confocal microscope (Leica).

### Flow Cytometer Analysis

WI-38 cells were incubated in 100 ng/ml (300 nM) DAPI-containing DMEM with or without 0.0085% SDS. After washing away, cells were fixed with 4% PFA and resuspended in flow cytometry buffer (1x PBS with 2% FBS). Flow cytometric analysis was performed by Amnis ImageStreamx Mk II (Millipore), and data were processed with FCS express (De novo software). Cell aggregates and debris were removed by standard gating strategy.

### Sample preparation for FIB-SEM

Normal human fibroblast cells (HCA-2) were fixed with a 3% glutaraldehyde solution in 100 mM cacodylate buffer (pH7.4) for 1 h at 4°C. The cells were then washed with 7.5% sucrose and then en bloc staining was performed following a procedure from NCMIR methods for 3D EM (https://ncmir.ucsd.edu/sbem-protocol). Epon 812-embedded specimens were glued onto an aluminum FIB-SEM rivet with conductive epoxy resin. 15 × 15 μm (1000 × 1000 pixels) images were acquired at 20 nm intervals. More than 200 images were collected to reconstitute a single cell.

### Stack alignment, segmentation, and 3D representation

The image stacks were automatically aligned by Fiji/ImageJ software. Plasma membrane structure segmentation and three-dimensional image-reconstitution were performed manually with the AMIRA software (FEI Visualization Science Group, Burlington, MA, USA).

### Calcium imaging

WI-38 cells (PDL: 43.7) were plated on glass-bottom dishes (CELLview™ Cell Culture Dish with Glass Bottom, Greiner). Cells loaded with a cytosolic Ca^2+^ indicator (5 μM Cal Red R525/650, AAT Bioquest) were imaged by a confocal microscope (SP8, Leica). Cytosolic Ca^2+^ dynamics were tracked with three stack images around the focal plane. Fluorescence intensity (Ex 492 nm/Em 500–550 nm and Ex 492 nm/Em 625–675 nm) were recorded every 15 seconds. For quantification, z stack images were prepared by Z projection tools (sum slices), and the whole area of each cell was selected by ROI selection tools (ImageJ, NIH). The average fluorescence ratio (500-550 nm/625-675 nm) was measured and calculated as F value. ΔF values were calculated from (F-F0) where F0 values were defined by averaging four frames before stimulation.

### RNA extraction and quantitative reverse-transcription polymerase chain reaction (qRT-PCR) analyses

Cells were lysed in TRIzol. RNA was extracted and purified using RNA Clean and Concentrator-5 kit (ZYMO Research) following the manufacturer’s instructions. Complementary DNA (cDNA) was synthesized using SuperScript™ IV VILO™ Master Mix with ezDNase enzyme kit (Thermo Fisher Scientific). The expression of target genes was determined using QuantStudio™ 1 Real-Time PCR system (Thermo Fisher Scientific). PCR amplification was performed using PowerTrack™ SYBR™ Green Master Mix (Thermo Fisher Scientific) with primer pairs as listed in Table S6. PCR was performed with the following thermocycling parameters; 2 min of initial DNA polymerase activation and DNA denaturation at 95°C, followed by 40 cycles of denaturation at 95°C for 5 sec, and primer annealing at 60°C for 30 sec. Melt curves of the qPCR products were analyzed from 65°C to 95°C. The data were statistically analyzed using SPSS software, version 21.0 (IBM Corp.). One-way analysis of variance (ANOVA) and Tukey-Kramer test were used to check the significant changes of gene expression. NormFinder software was used to determine the stability values of the reference gene. Changes in gene expression, expressed as fold change, were calculated using the ΔΔCt method, where Actb was used as a reference gene for normalizing the expression. The nested scatter plot was generated to show the fold changes of each replicate (GraphPad Prism).

### EdU staining

Young WI-38 cells (PDL32-46) were used as controls. For replication-induced senescence, WI-38 were passaged until they lost the ability to proliferate and became fully senescent around PDL52. For the plasma membrane damage-dependent senescence and DNA damage-dependent senescence, Young WI-38 cells (PDL32-46) were treated with 0.0090% SDS or 250 nM doxorubicin (Cayman Chemical) containing medium for 24 hours. The cells were washed twice with fresh medium, then cultured for 16 days with changing medium every four days. Click-iT Plus EdU Cell Proliferation Kit for Imaging (Thermo Fisher Scientific) with Alexa Fluor 647 dye was used for the detection of DNA synthesis. Cells were labeled with 10 μM EdU for 24 hours, then treated according to the manufacturer’s instructions. EdU-coupled Alexa Fluor 647 signals and counter-stained Hoechst 33342 signals were obtained with AxioObserver7 (Zeiss).

## General

We are grateful to T. Hunt, D. Pellman, Z. Storchová, I. Fukunaga, A. Nishiyama, M. Shimada, T. Kiyomitsu, R. Hatakeyama, and S. Yoshida for critical reading of the manuscript; Y. Hamamura for live-imaging analysis, S. Arai, and S. Enomoto for FIB-SEM analysis; H.I. Wen, T. Inden, C. Yamada-Namikawa, K. Mori, M. Akiyama, Y. Tsukui, Y. Matsui, K. Koizumi, T. Mochizuki, S. Komoto and P. Barzaghi for technical assistance. A portion of this work was conducted in the Institute of Transformative Bio-Molecules (WPI-ITbM) at Nagoya University, supported by the Japan Advanced Plant Science Network. A portion of this work was supported by NIMS microstructural characterization platform as a program of “Nanotechnology Platform” of MEXT Japan. A portion of this work was performed with the help of OIST imaging section members.

## Author contributions

K.K. and M.N. planned the overall studies and interpreted the data; K.K., and S.S. performed yeast experiments; K.K., Y.J., Y. Moriyama., K.S., N.B.R., Y.C., K.N., Y. Masukagami, and H.B. performed human cell culture experiments; H.T. performed and interpreted yeast electron microscope experiments; Y. Moriyama performed and interpreted FIB-SEM experiments, K.K., Y.S. and T.H. performed and interpreted laser damage experiments; K.K., Y.J., Y. Moriyama., K.S., N.B.R., Y.C., K.N. and M.N. prepared the Figures; K.K. and M.N. wrote the manuscript with editing by all authors.

## Data and materials availability

All raw data are available upon request.

---

**Fig. S1.**
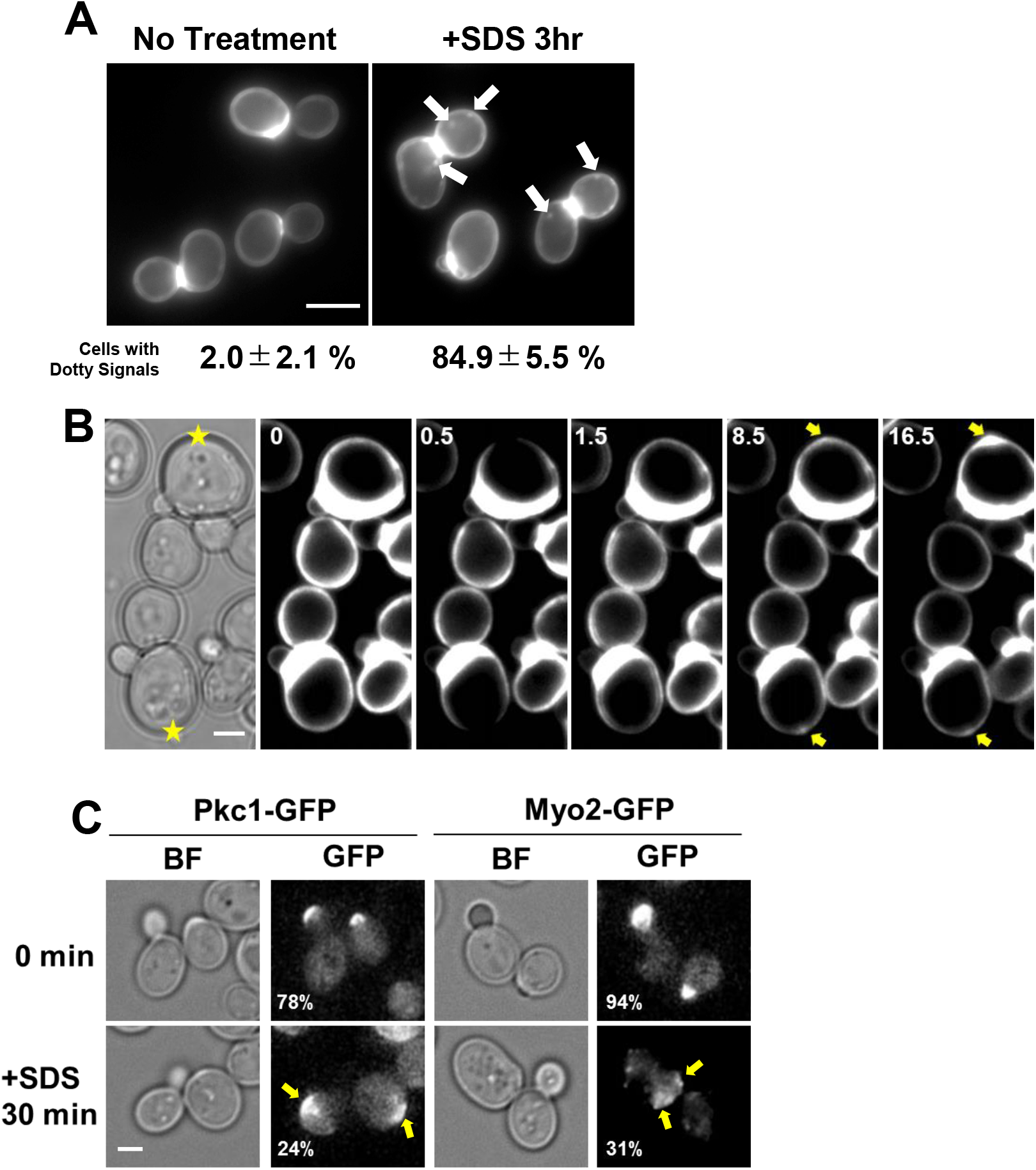

**Fig. S3.**
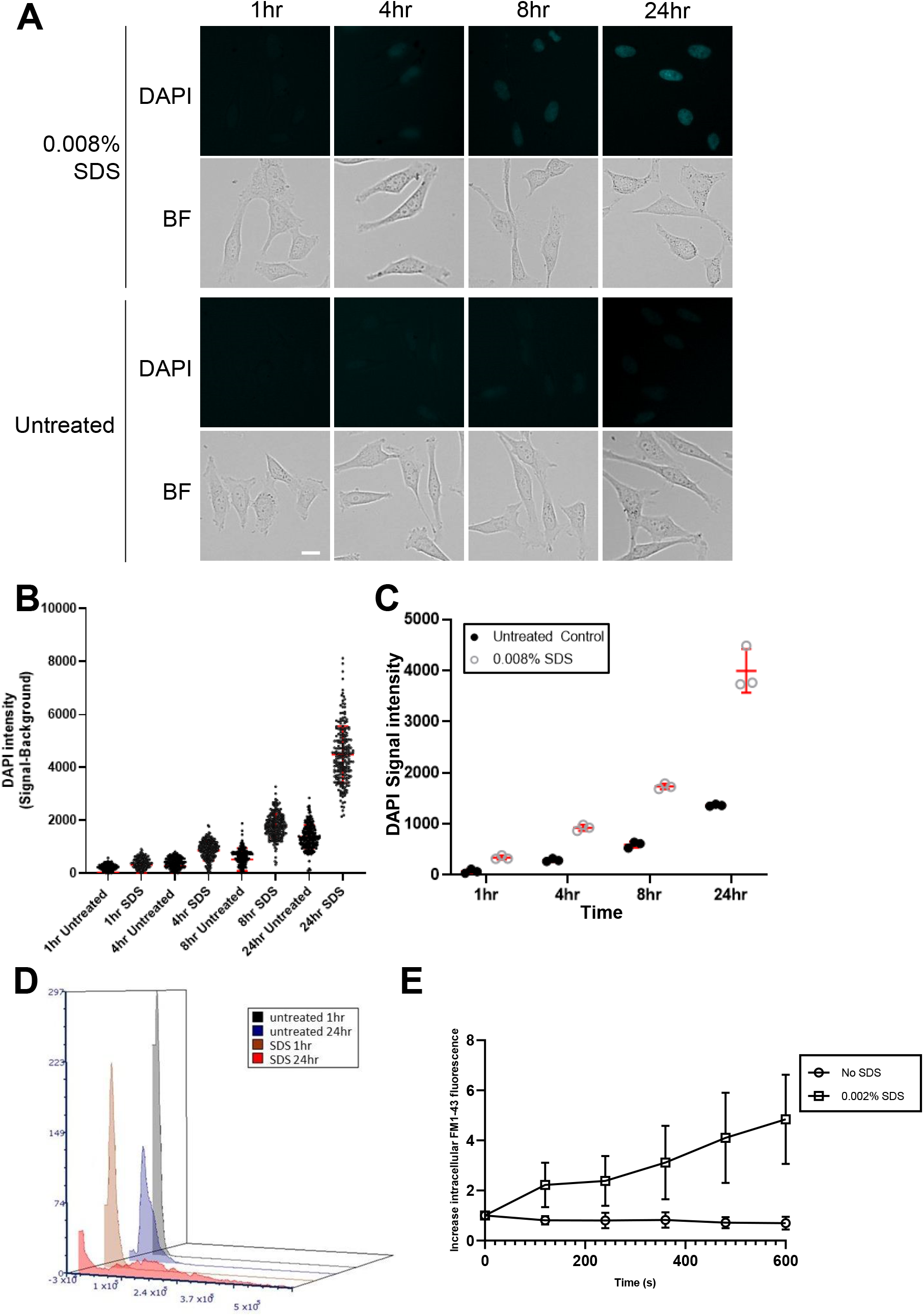

**Fig. S3.**
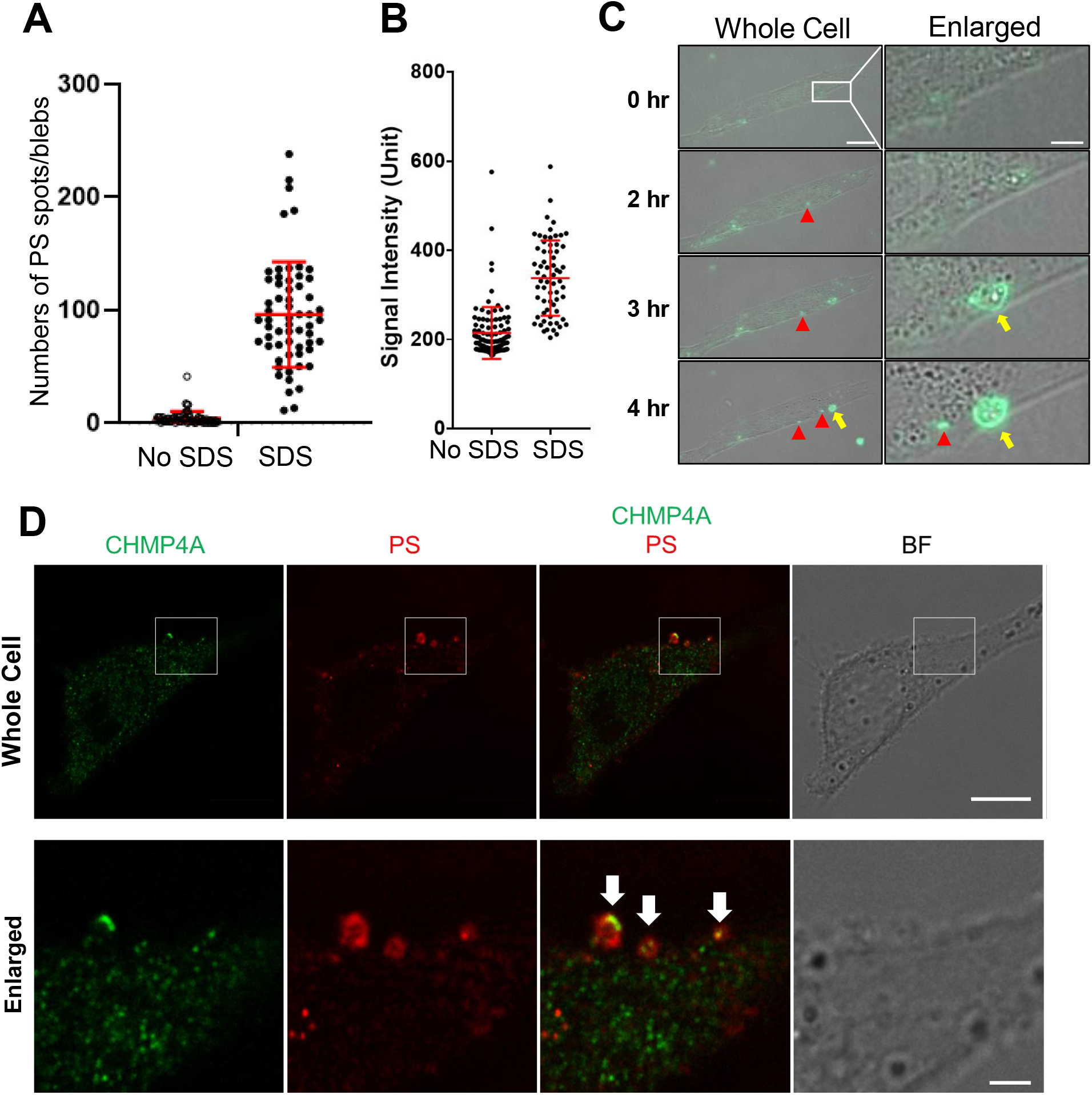

**Fig. S4.**
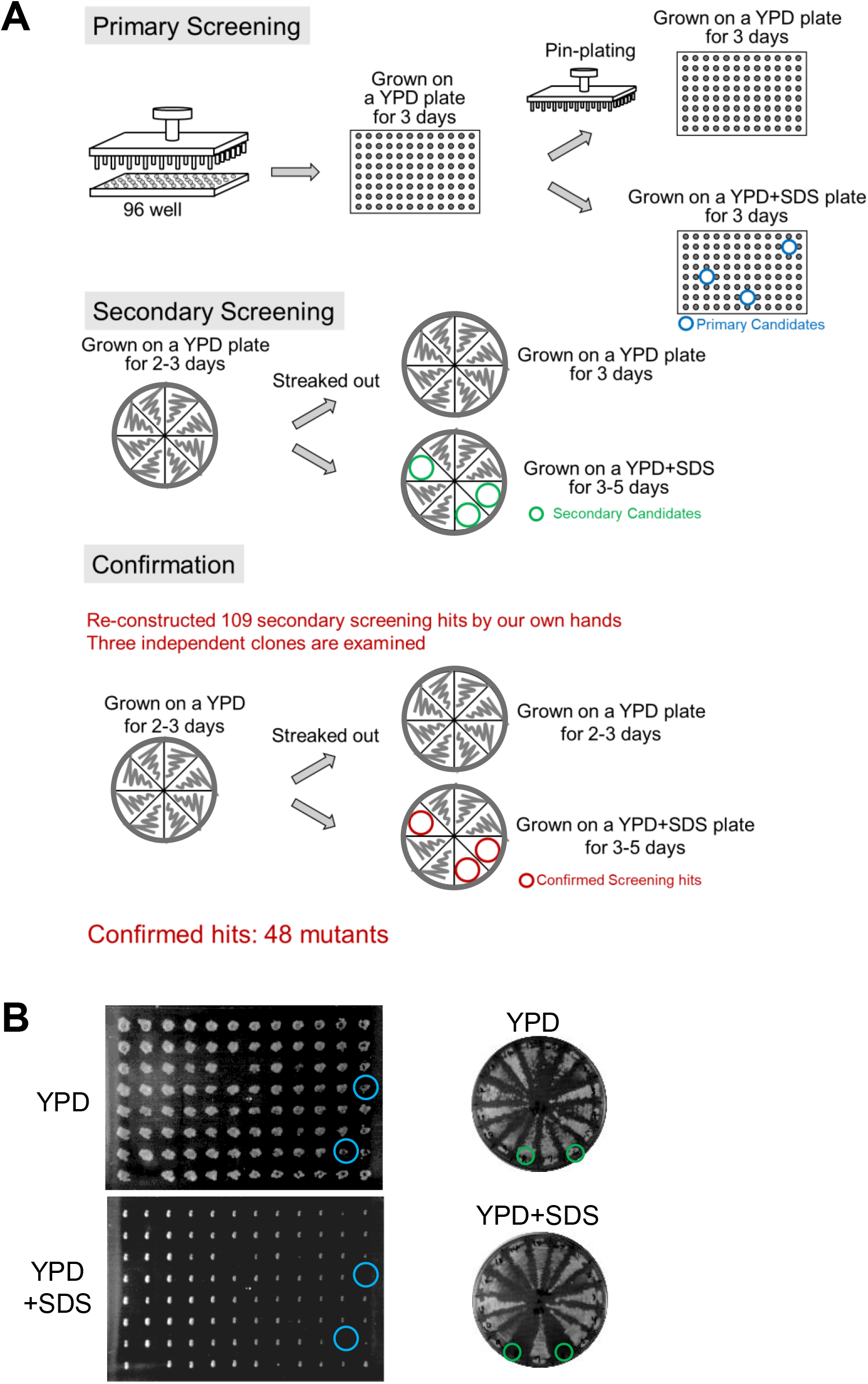

**Fig. S5.**
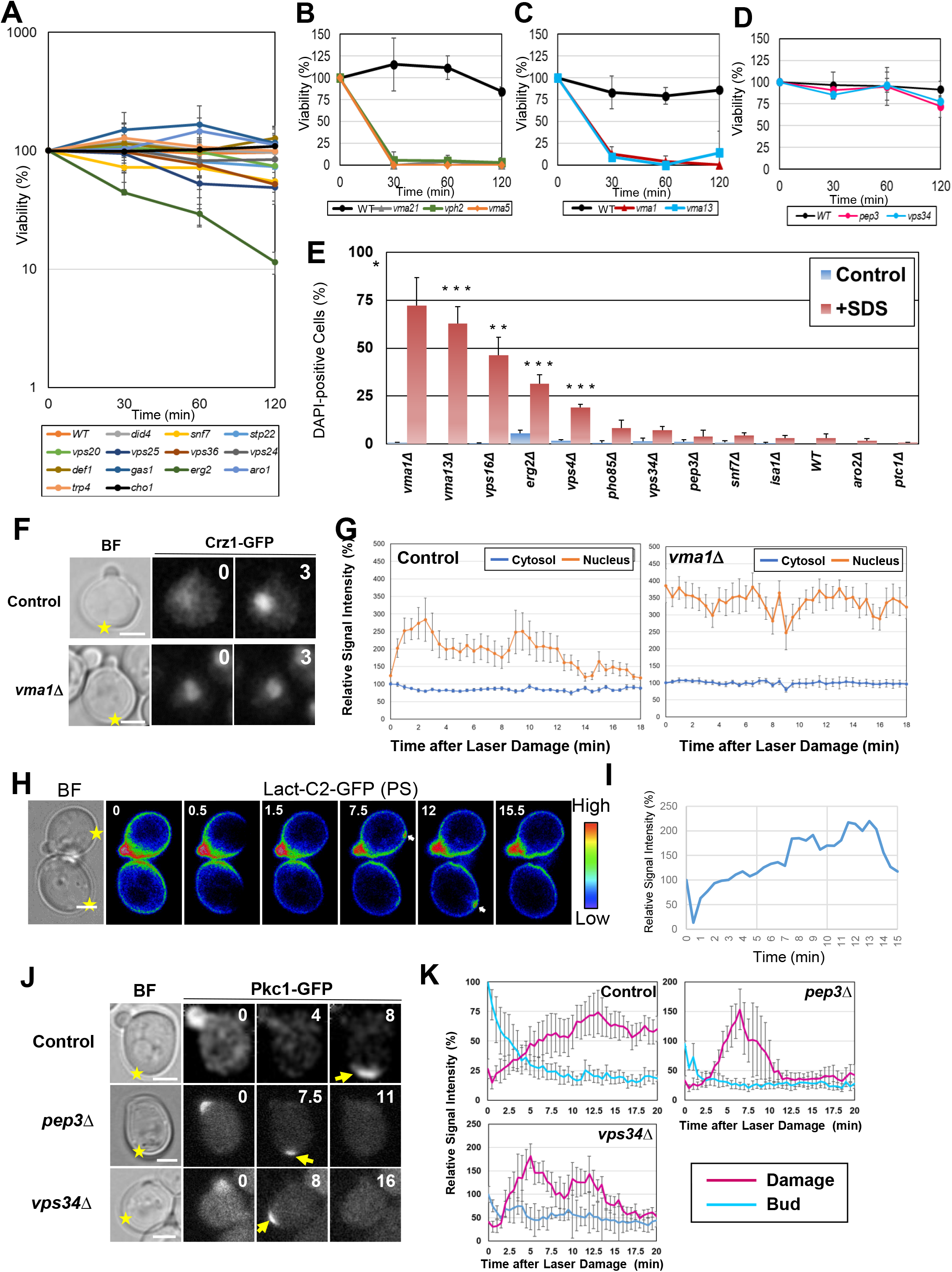

**Fig. S6.**
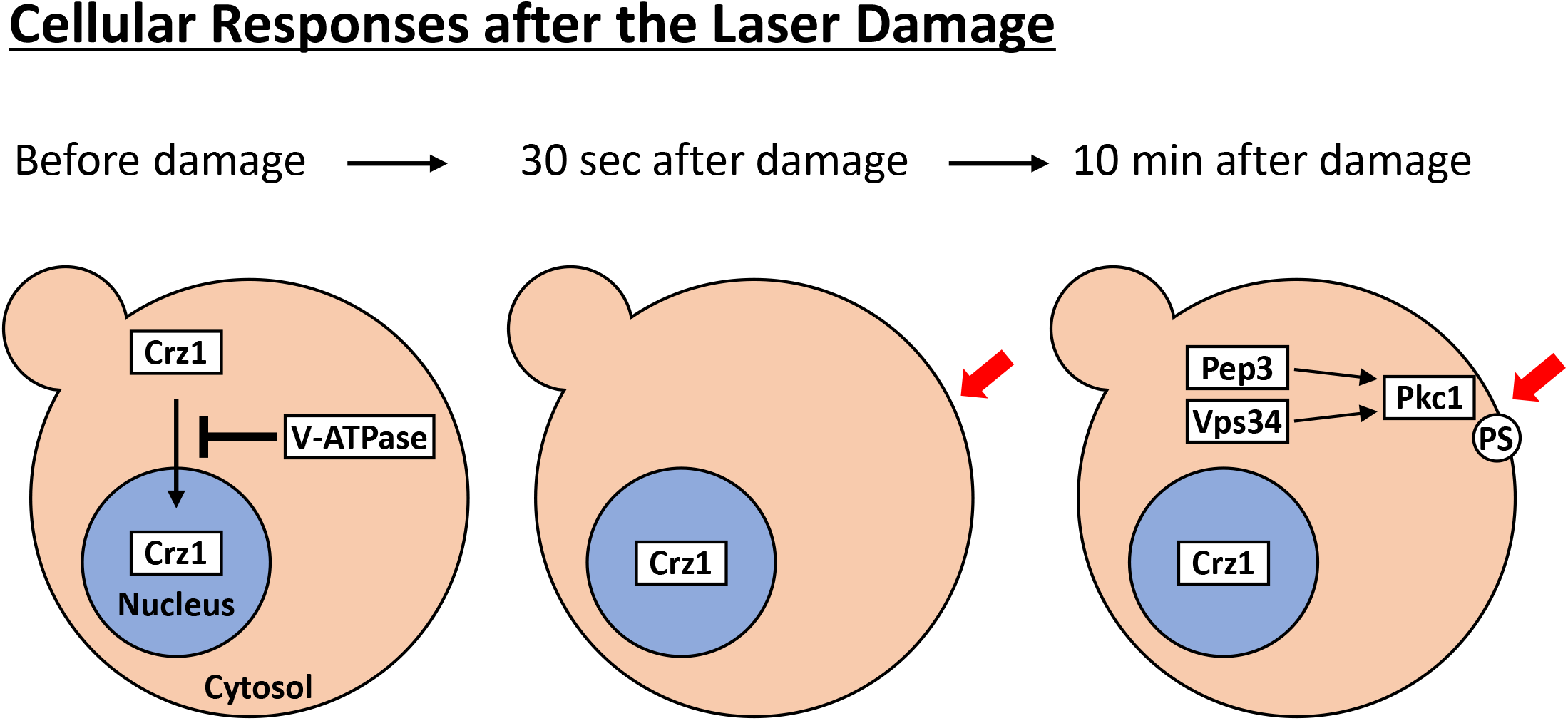

**Fig. S7.**
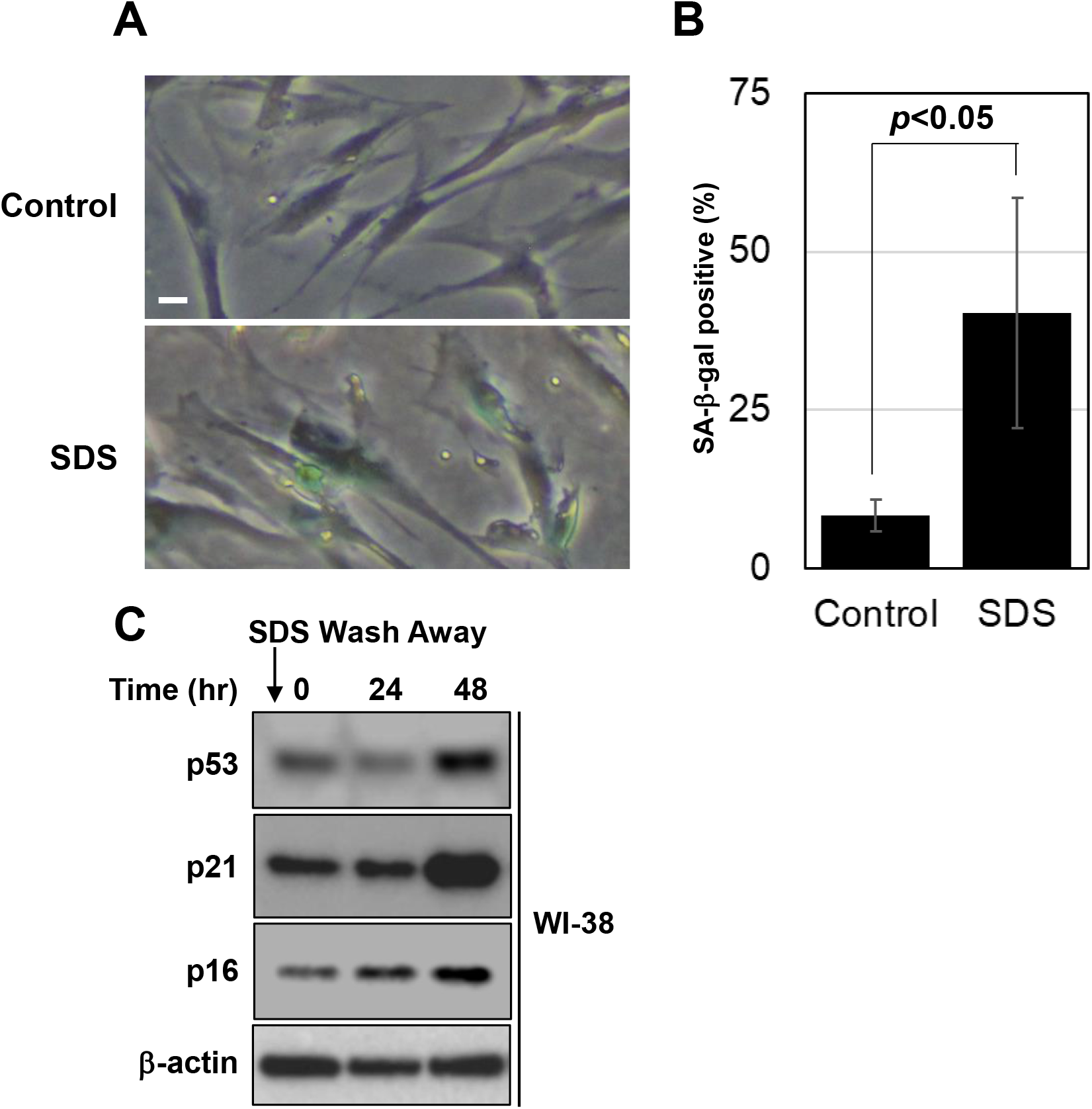

**Fig. S8.**
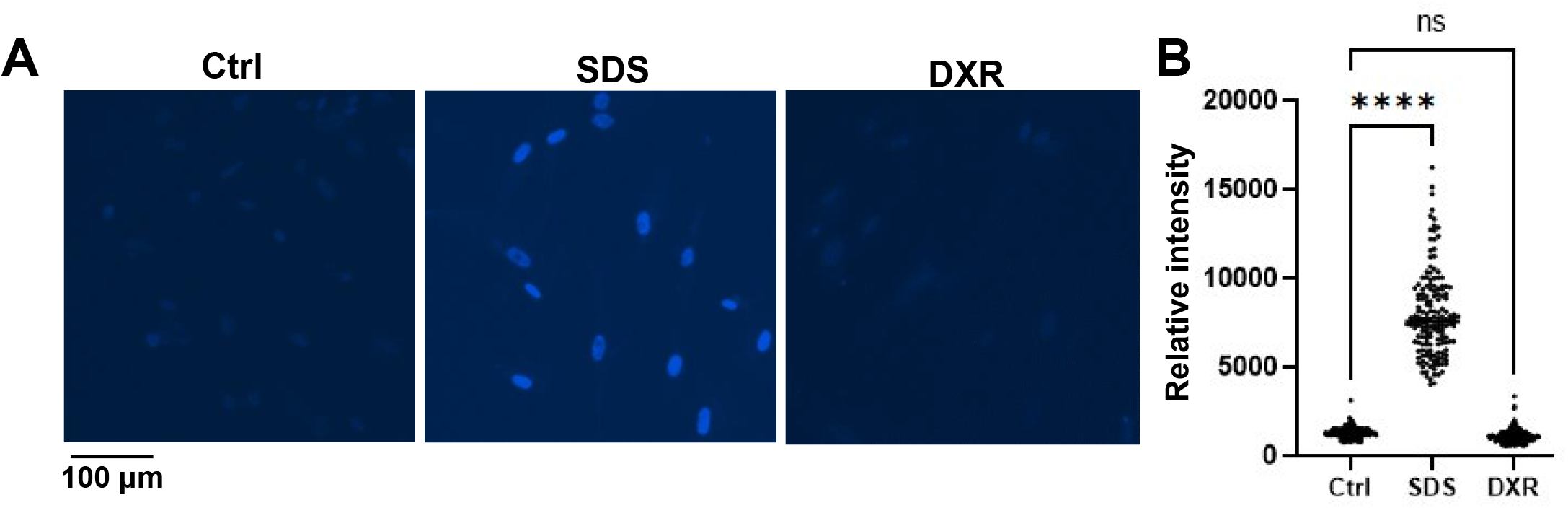

**Table S1.**
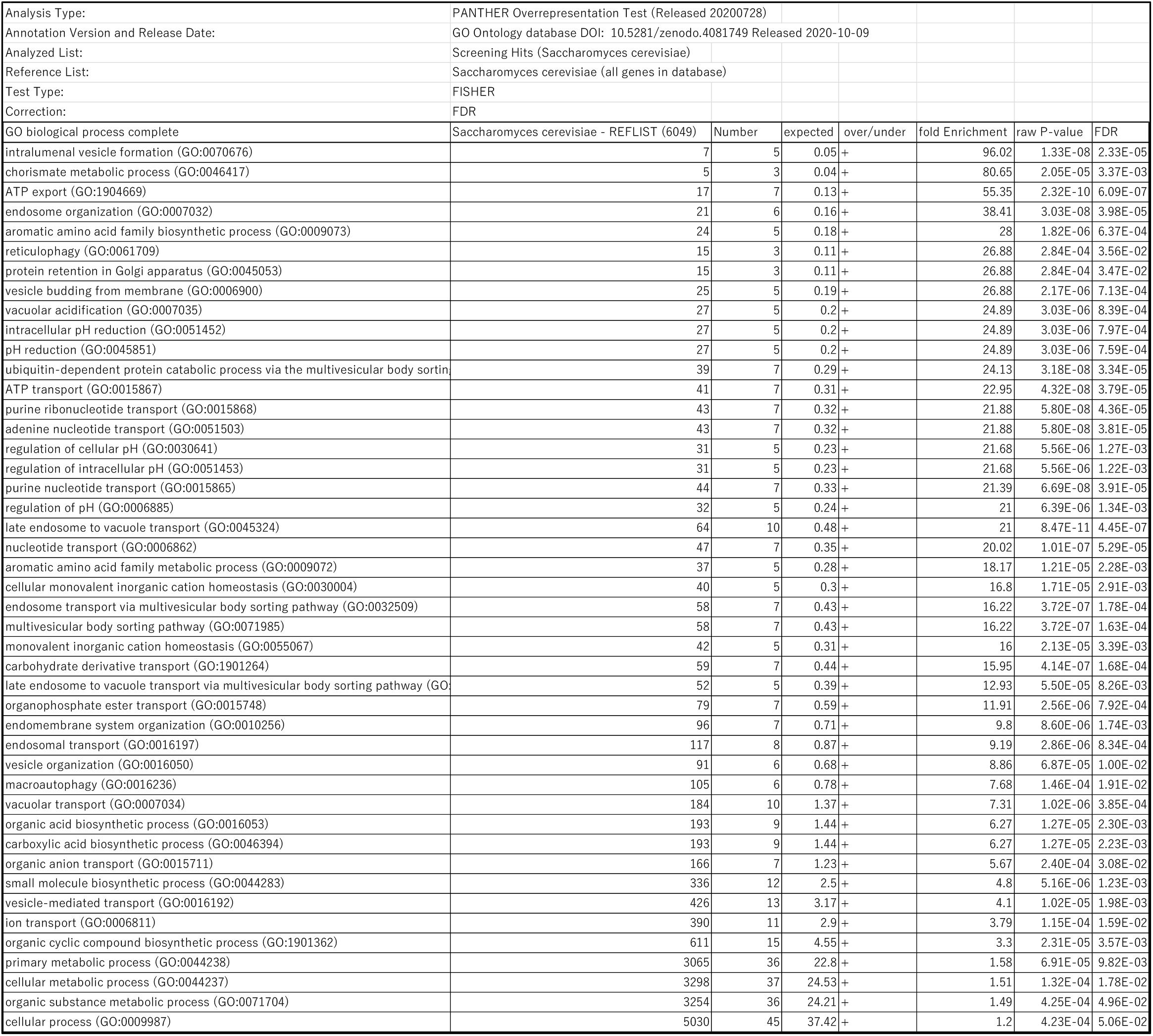

**Table S2.**
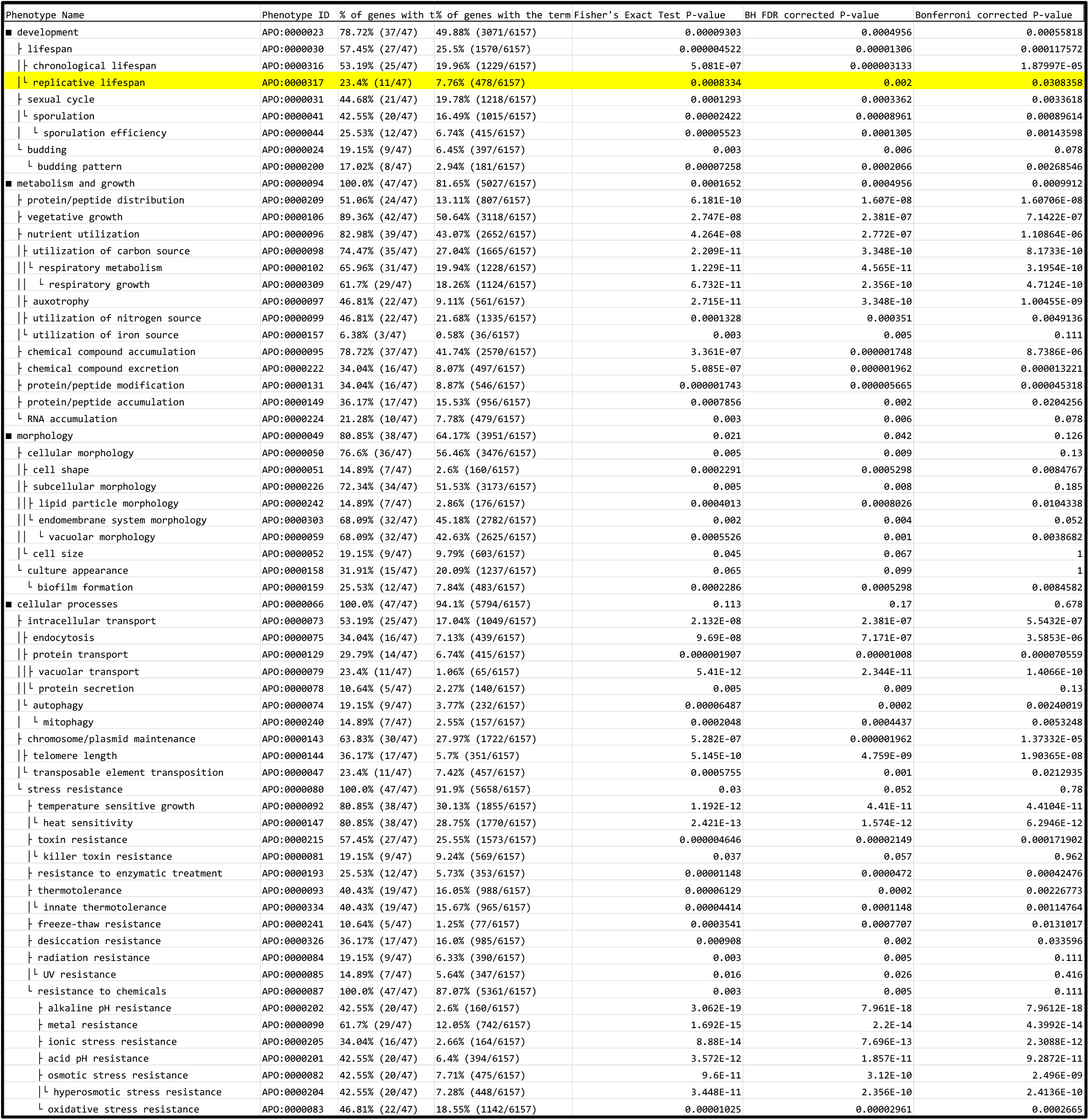

**Table S3.**
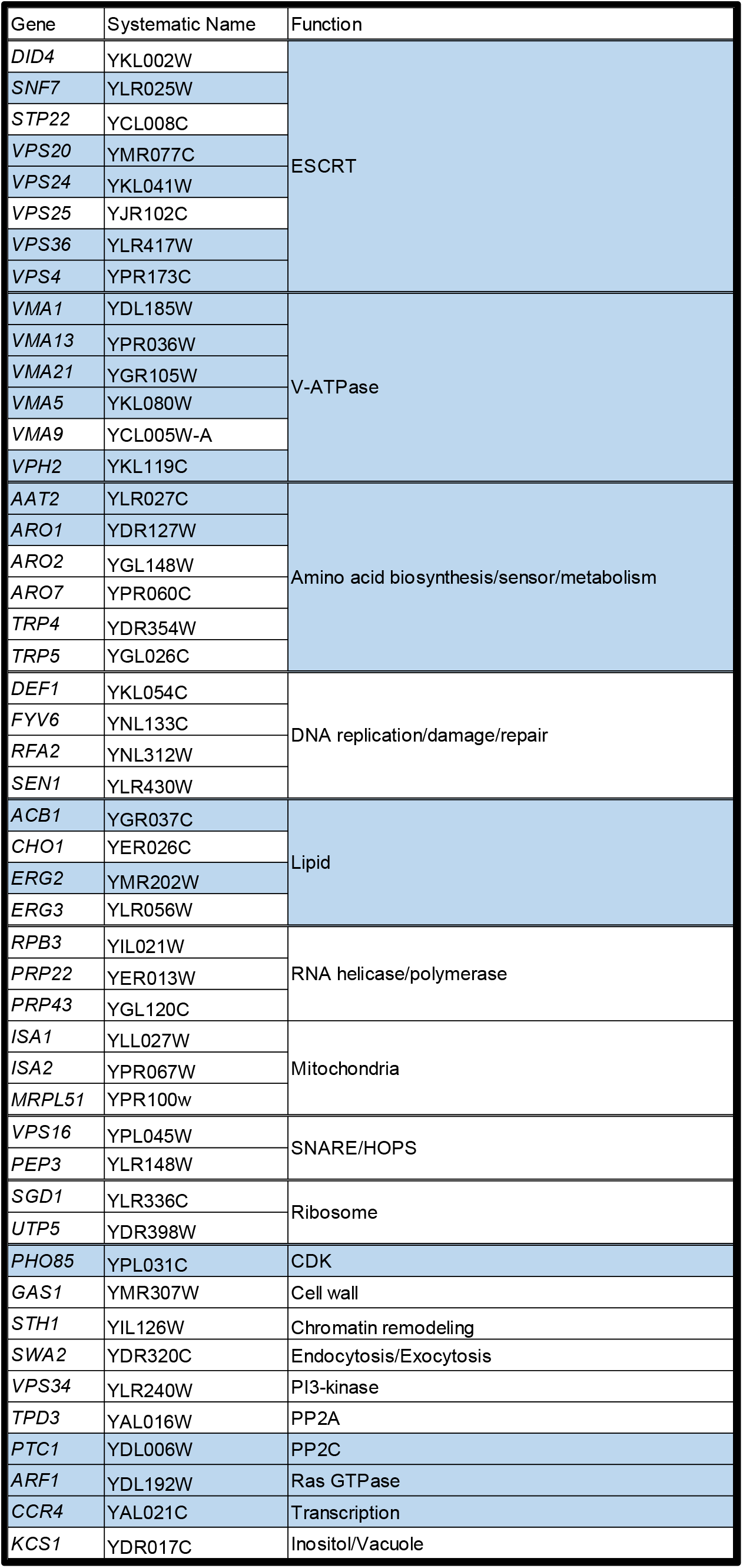
The genes and gene functions associated with “replicative lifespan” are highlighted in blue.

**Table S4.**

| Strain Name | Alias | MAT | Genotype | Source |
| --- | --- | --- | --- | --- |
| YKK1471 | BY23849 | a | *his3, leu2, met15, ura3* | National Bio-Resource Project (NBRP) of the MEXT, Japan |
| YKK107 | PY6767 | a | *PKC1::GFP::HIS3* | Kono *et al.*, 2012 |
| YKK2969 |  | a | *MYO2::GFP::HIS3* | Yeast GFP clone collection (ThermoFisher) |
| YKK847 |  | a | *CRZ1::GFP::HIS3* | Yeast GFP clone collection (ThermoFisher) |
| YKK1488 |  | a | *vma1::Km CRZ1-GFP:HIS3* | This study |
| YKK1550 |  | a | *pep3::Km PKC1-GFP:HIS3* | This study |
| YKK1516 |  | a | *vps34::Km PKC1-GFP:HIS3* | This study |
| YKK1591 |  | a | YKK1471 (pRH176) | This study |
| YKK2407 |  | a | *Km::pTEF1::VPS4* | This study |
| Yeast deletion collection (Winzeler et al., 1999) |  |  |  |  |
| Yeast DAmP library (Breslow et al., 2008) |  |  |  |  |

| Plasmid Name | Description | Source |
| --- | --- | --- |
| pRH176 | *CEN, URA3, PGPD-GFP-LactC2* | Riko Hatakeyama & Satoshi Yoshida |

**Table S5.**

| Antibodies | Catalog No | Species | Source |
| --- | --- | --- | --- |
| Anti-CHMP4A | HPA06873 | Rabbit | Sigma-Aldrich (Merck), St. Luis, MO |
| Anti-p53 [DO-1] | sc-126 | Mouse | Santa Cruz Biotechnologies, Santa Cruz, CA |
| Anti-p21 Waf1/Cip1 [F-5] | sc-6246 | Mouse | Santa Cruz Biotechnologies, Santa Cruz, CA |
| Anti-p21 [EPR362] | ab109520 | Rabbit | Abcam, Cambridge, UK |
| Anti-p16 (JC8) | sc-56330 | Mouse | Santa Cruz Biotechnologies, Santa Cruz, CA |
| Anti-CDKN2A/p16INK4a [EPR1473] | ab108349 | Rabbit | Abcam, Cambridge, UK |
| Anti-Phospho-ATM (S1981) | AAF1655 | Rabbit | R&D Systems, Minneapolis, MN |
| Anti-ATM [2C1] | sc-23921 | Mouse | Santa Cruz Biotechnologies, Santa Cruz, CA |
| Anti-Phospho-H2A.X(Ser139) [JBW301] | 05-636 | Mouse | Sigma-Aldrich (Merck), St. Luis, MO |
| Anti-H2A.X | ab11175 | Rabbit | Abcam, Cambridge, UK |
| Anti-Phospho-Chk1(Ser345) [133D3] | #2348 | Rabbit | Cell Signaling Technology, Boston, MA |
| Anti-Chk1 [DCS310] | C9358 | Mouse | Sigma-Aldrich (Merck), St. Luis, MO |
| Anti-GFP | A-6455 | Rabbit | Thermo Fisher Scientific, Waltham, MA |
| Anti-GAPDH (D16H11) | 5174 | Rabbit | Cell Signaling Technology, Boston, MA |
| Anti-beta Actin antibody [AC-15] | ab6276 | Mouse | Abcam, Cambridge, UK |

**Table S6.**
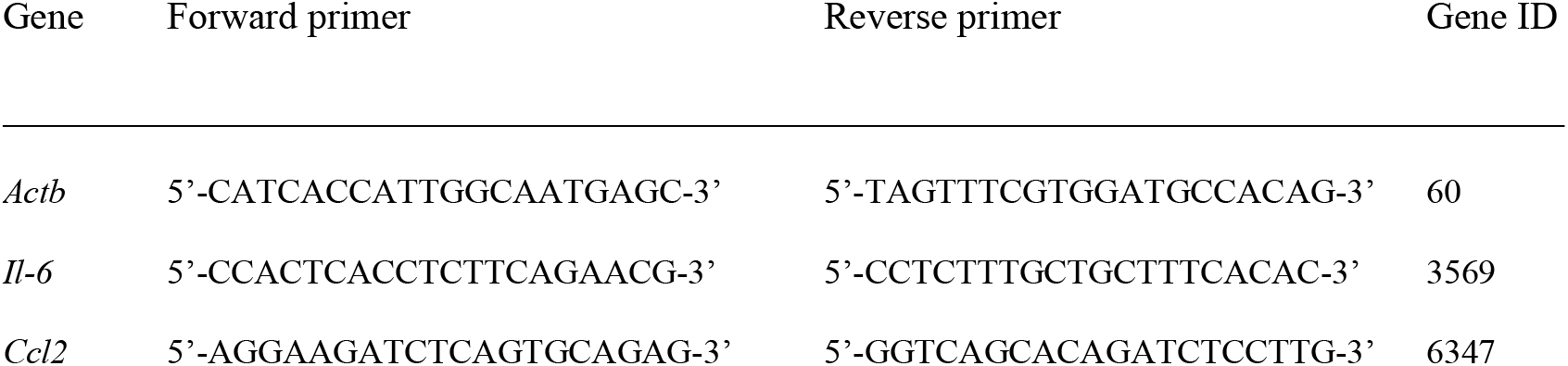

## References

1. N.W. Andrews, P.E. Almeida, M. Corrotte, Damage control: cellular mechanisms of plasma membrane repair. Trends. Cell Biol. 24, 734–742 (2014).

2. P.L. McNeil, R.A. Steinhardt, Plasma membrane disruption: repair, prevention, adaptation. Annu. Rev. Cell Dev. Biol. 19, 697–731 (2003).

3. K.J. Sonnemann, W.M. Bement, Wound repair: toward understanding and integration of single-cell and multicellular wound responses. Annu. Rev. Cell Dev. Biol. 27, 237–263 (2011).

4. R. Bashir, S. Britton, T. Strachan, S. Keers, E. Vafiadaki, M. Lako, I. Richard, S. Marchand, N. Bourg, Z. Argov, M. Sadeh, I. Mahjneh, G. Marconi, M.R. Passos-Bueno, E. de S. Moreira, M. Zatz, J.S. Beckmann, K. Bushby, A gene related to Caenorhabditis elegans spermatogenesis factor fer-1 is mutated in limb-girdle muscular dystrophy type 2B. Nat. Genet. 20, 37–42 (1998).

5. J. Suzuki, M. Umeda, P.J. Sims, S. Nagata, Calcium-dependent phospholipid scrambling by TMEM16F. Nature. 468, 834–838 (2010).

6. N. Wu, V. Cernysiov, D. Davidson, H. Song, J. Tang, S. Luo, Y. Lu, J. Qian, I.E. Gyurova, S.N. Waggoner, V.Q. Trinh, R. Cayrol, A. Sugiura, H.M. McBride, J.F. Daudelin, N. Labrecque, A. Veillette, Critical Role of Lipid Scramblase TMEM16F in Phosphatidylserine Exposure and Repair of Plasma Membrane after Pore Formation. Cell Rep. 30, 1129–1140.e5 (2020).

7. P.L. McNeil, T. Kirchhausen, An emergency response team for membrane repair. Nat. Rev. Mol. Cell Biol. 6, 499–505 (2005).

8. M. Nakamura, A.N.M. Dominguez, J.R. Decker, A.J. Hull, J.M. Verboon, S.M. Parkhurst, Into the breach: how cells cope with wounds. Open Biol. 8, 180135 (2018).

9. Y. Zhen, M. Radulovic, M. Vietri, H. Stenmark. Sealing holes in cellular membranes. EMBO J. e106922 (2021).

10. B.M. Burkel, H.A. Benink, E.M. Vaughan, G. von Dassow, W.M. Bement, A Rho GTPase signal treadmill backs a contractile array. Dev. Cell 23, 384–396 (2012).

11. M. Vietri, M. Radulovic, H. Stenmark, The many functions of ESCRTs. Nat. Rev. Mol. Cell Biol. 21, 25–42 (2020).

12. B.T. Cookson, M.A. Brennan, Pro-inflammatory programmed cell death, Trends Microbiol. 9, 113–114 (2001).

13. L. Wang, X. Qin, J. Liang, P. Ge, Induction of Pyroptosis: A Promising Strategy for Cancer Treatment. Front Oncol. 11, 635774 (2021).

14. K. Kono, Y. Saeki, S. Yoshida, K. Tanaka, D. Pellman, Proteasomal degradation resolves competition between cell polarization and cellular wound healing. Cell 150, 151–164 (2012).

15. K. Kono, A. Al-Zain, L. Schroeder, M. Nakanishi, A.E. Ikui, Plasma membrane/cell wall perturbation activates a novel cell cycle checkpoint during G1 in Saccharomyces cerevisiae. Proc. Natl. Acad. Sci. USA. 113, 6910–6915 (2016).

16. M. Demaria, N. Ohtani, S.A. Youssef, F. Rodier, W. Toussaint, J.R. Mitchell, R.M. Laberge, J. Vijg, H. Van Steeg, M.E. Dollé, J.H. Hoeijmakers, A. de Bruin, E. Hara, J. Campisi. An essential role for senescent cells in optimal wound healing through secretion of PDGF-AA. Dev Cell. 31, 722–33 (2014).

17. A. Draeger, K. Monastyrskaya, E.B. Babiychuk, Plasma membrane repair and cellular damage control: the annexin survival kit. Biochem Pharmacol. 81, 703–712 (2011).

18. E.A. Winzeler, D.D. Shoemaker, A. Astromoff, H. Liang, K. Anderson, B. Andre, R. Bangham, R. Benito, J.D. Boeke, H. Bussey, A.M. Chu, C. Connelly, K. Davis, F. Dietrich, S.W. Dow, M.E. Bakkoury, F. Foury, S.H. Friend, E. Gentalen, G. Giaever, J.H. Hegemann, T. Jones, M. Laub, H. Liao, N. Liebundguth, D.J. Lockhart, A. Lucau-Danila, M. Lussier, N. M’Rabet, P. Menard, M. Mittmann, C. Pai, C. Rebischung, J.L. Revuelta, L. Riles, C.J. Roberts, P. Ross-MacDonald, B. Scherens, M. Snyder, S. Sookhai-Mahadeo, R.K. Storms, S. Véronneau, M. Voet, G. Volckaert, T.R. Ward, R. Wysocki, G.S. Yen, K. Yu, K. Zimmermann, P. Philippsen, M. Johnston, R.W. Davis, Functional characterization of the S. cerevisiae genome by gene deletion and parallel analysis. Science 285, 901–906 (1999).

19. D.K. Breslow, D.M. Cameron, S.R. Collins, M. Schuldiner, J. Stewart-Ornstein, H.W. Newman, S. Braun, H.D. Madhani, N.J. Krogan, J.S. Weissman, A comprehensive strategy enabling high-resolution functional analysis of the yeast genome. Nat. Methods 5, 711–718 (2008).

20. M.P. Weng, B.Y. Liao, modPhEA: model organism Phenotype Enrichment Analysis on eukaryotic gene sets. Bioinformatics 33, 3505–3507 (2017).

21. D.A. Sinclair, Studying the replicative life span of yeast cells. Methods Mol. Biol. 1048, 49–63 (2013).

22. Y. Johmura, M. Shimada, T. Misaki, A. Naiki-Ito, H. Miyoshi, N. Motoyama, N. Ohtani, E. Hara, M. Nakamura, A. Morita, S. Takahashi, M. Nakanishi, Necessary and sufficient role for a mitosis skip in senescence induction. Mol. Cell 55, 73–84 (2014).

23. A. Rufini, P. Tucci, I. Celardo, G. Melino, Senescence and aging: the critical roles of p53. Oncogene 32, 5129–5143 (2013).

24. F. d’Adda di Fagagna, P.M. Reaper, L. Clay-Farrace, H. Fiegler, P. Carr, T. Von Zglinicki, G. Saretzki, N.P. Carter, S.P. Jackson, A DNA damage checkpoint response in telomere-initiated senescence. Nature 426, 194–198 (2003).

25. T.D. Halazonetis, V.G. Gorgoulis, J. Bartek, An oncogene-induced DNA damage model for cancer developlasma membraneent. Science 319, 1352–1355 (2008).

26. N. Martin, D. Bernard, Calcium signaling and cellular senescence. Cell Calcium 70, 16–23 (2018).

27. J.A. Schäfer, J.P. Schessner, P.W. Bircham, T. Tsuji, C. Funaya, O. Pajonk, K. Schaeff, G. Ruffini, D. Papagiannidis, M. Knop, T. Fujimoto, S. Schuck, ESCRT machinery mediates selective microautophagy of endoplasmic reticulum in yeast. EMBO J. 39, e102586 (2020).

28. A.J. Jimenez, P. Maiuri, J. Lafaurie-Janvore, S. Divoux, M. Piel, F. Perez, ESCRT machinery is required for plasma membrane repair. Science 343, 1247136 (2014).

29. O. Schmidt, Y. Weyer, S. Sprenger, M.A. Widerin, S. Eising, V. Baumann, M. Angelova, R. Loewith, C.J. Stefan, M.W, Hess, F. Fröhlich, D. Teis, TOR complex 2 (TORC2) signaling and the ESCRT machinery cooperate in the protection of plasma membrane integrity in yeast. J Biol Chem, 295, 12028–12044 (2020).

30. A.L. Hughes, Gottschling DE, An early age increase in vacuolar pH limits mitochondrial function and lifespan in yeast. Nature. 492, 261–265 (2012).

31. C.E. Hughes, T.K. Coody, M.Y. Jeong, J.A. Berg, D.R. Winge, A.L. Hughes, Cysteine Toxicity Drives Age-Related Mitochondrial Decline by Altering Iron Homeostasis. Cell. 180, 296–310.e18 (2020).

32. V. Lundblad, J.W. Szostak, A mutant with a defect in telomere elongation leads to senescence in yeast. Cell. 57, 633–643 (1989).

33. E. Henninger, M.T. Teixeira, Telomere-driven mutational processes in yeast. Curr Opin Genet Dev. 60, 99–106 (2020).

34. R.G. Syljuåsen, S. Jensen, J. Bartek, J. Lukas. Adaptation to the ionizing radiation-induced G2 checkpoint occurs in human cells and depends on checkpoint kinase 1 and Polo-like kinase 1 kinases. Cancer Res. 66, 10253–10257 (2006).

35. H. Coutelier, Z. Xu, M.C. Morisse, M. Lhuillier-Akakpo, S. Pelet, G. Charvin, K. Dubrana, M.T. Teixeira, Adaptation to DNA damage checkpoint in senescent telomerase-negative cells promotes genome instability. Genes Dev. 32, 1499–1513 (2018).

36. T. Arnould, S. Vankoningsloo, P. Renard, A. Houbion, N. Ninane, C. Demazy, J. Remacle, M. Raes, CREB activation induced by mitochondrial dysfunction is a new signaling pathway that impairs cell proliferation. EMBO J. 1, 53–63 (2002).

37. S. Nagata, T. Sakuragi, K. Segawa, Flippase and scramblase for phosphatidylserine exposure. Curr Opin Immunol. 62, 31–38 (2020).

38. Y. Johmura, T. Yamanaka, S. Omori, T.W. Wang, Y. Sugiura, M. Matsumoto, N. Suzuki, S. Kumamoto, K. Yamaguchi, S. Hatakeyama, T. Takami, R. Yamaguchi, E. Shimizu, K. Ikeda, N. Okahashi, R. Mikawa, M. Suematsu, M. Arita, M. Sugimoto, K.I. Nakayama, Y. Furukawa, S. Imoto, M. Nakanishi. Senolysis by glutaminolysis inhibition ameliorates various age-associated disorders. Science. 371, 265–270 (2021).

39. M. Ge, L. Hu, H. Ao, M. Zi, Q. Kong, Y. He, Senolytic targets and new strategies for clearing senescent cells. Mech Ageing Dev. 111468 (2021).

40. J. Schindelin, I. Arganda-Carreras, E. Frise, V. Kaynig, M. Longair, T. Pietzsch, S. Preibisch, C. Rueden, S. Saalfeld, B. Schmid, J.Y. Tinevez, D.J. White, V. Hartenstein, K. Eliceiri, P. Tomancak, A. Cardona, Fiji: an open-source platform for biological-image analysis. Nat. Methods 9, 676–682 (2012).

41. F. Madeo, E. Frohlich, K.U. Frohlich, A yeast mutant showing diagnostic markers of early and late apoptosis. J. Cell Biol. 139, 729–734 (1997).

42. G.P. Dimri, X. Lee, G. Basile, M. Acosta, G. Scott, C. Roskelley, E.E. Medrano, M. Linskens, I. Rubelj, O. Pereira-Smith, A biomarker that identifies senescent human cells in culture and in aging skin in vivo. Proc. Natl. Acad. Sci. USA 92, 9363–9367 (1995).

