## Supplemental Text 1 for "Plasma membrane damage limits replicative lifespan in yeast and induces premature senescence in human fibroblasts"

Keiko Kono et al.

**Supplementary Text**

To characterize the screening hits, we examined cell viability after SDS treatment. We found that ESCRT mutants (*did4∆*, *snf7∆*, *stp22∆*, *vps20∆*, *vps25∆*, *vps36∆*, and *vps24∆*) survived in the course of the two-hour experiment, whereas V-ATPase mutants (*vma21∆*, *vph2∆*, *vma5∆*, *vma1∆*, and *vma13∆*) lost their viability within 30 min (Fig. S5A-D).

V-ATPase produces a proton gradient across the vacuolar membrane, enabling Ca^2+^ uptake into the vacuole; the mutants lucking functional V-ATPase show high cytoplasmic Ca^2+^ levels (42). Since Ca^2+^ influx at the damage site is essential for membrane resealing in higher eukaryotes (6,7), SDS sensitivity in V-ATPase mutants may be explained by the high cytosolic Ca^2+^ concentration in V-ATPase mutants, preventing membrane resealing. Indeed, in our DAPI penetration assay (30 min incubation in YPD+SDS and quick wash with YPD, followed by 5 min incubation with DAPI), V-ATPase mutants (*vma1∆* and *vma13∆*) showed high DAPI-positivity (*vma1∆*: 72.2±14.6, *vma13∆*: 63.0±8.6; Fig. S5E) analogous to SDS and EGTA-treated wild type (Fig. 1A and B). These results raise a possibility that Ca^2+^ influx-dependent membrane resealing is impaired in V-ATPase mutants.

To test the possibility that Ca^2+^ homeostasis is defective in V-ATPase mutants, we monitored subcellular localization of Crz1-GFP, a Ca^2+^-responsive transcription factor that enters nucleus after various stimuli (43). After the laser-induced cell wall and plasma membrane damage, Crz1-GFP entered the nucleus within 30 sec (Fig. S5F and G). In a V-ATPase mutant *vma1∆*, even before the laser damage Crz1-GFP signals at the nucleus was comparable to the peak levels of control cells and did not increase after the laser damage. These results support our interpretation that Ca^2+^-influx detection is impaired in *vma1∆*.

Since *cho1∆*, defective in phospholipid phosphatidylserine (PS) synthesis, was a screening hit, we examined whether PS is involved in plasma membrane/cell wall repair processes after the laser damage. Wild type yeast cells harboring a plasmid of Lact-C2-GFP, a PS marker, were subjected to the laser damage assay. We found that Lact-C2-GFP signal gradually accumulated at the damage site and peaked after ~13 min (Fig. S5H and I), consistent with the idea that PS is involved in the plasma membrane/cell wall repair processes.

We next focused on *pep3*∆ and *vps34*∆. These mutants shared three common phenotypes: 1) the cell viability did not decrease after 2hr SDS treatment (Fig. S5D), 2) the cells did not show DAPI penetration after 30 min SDS treatment (Fig. S5E), and 3) in the laser damage experiment, a major repair protein Pkc1-GFP failed to be retained at the laser damage site (Fig. S5J and K). These results suggest that Pep3 and Vps34 are required for the retention of Pkc1 but not for plasma membrane resealing immediately after the damage or initial Pkc1 recruitment to the damage site.

In summary, here we revealed four cellular processes during plasma membrane/cell wall damage response in budding yeast: 1) V-ATPase-dependent prevention of immediate cell death, 2) Crz1 nuclear import, 3) PS recruitment to the damage site, and 4) Pep3 and Vps34-dependent retention of Pkc1 (Fig. S6).

**Supplementary Reference**

42. Y. Ohya, N. Umemoto, I. Tanida, A. Ohta, H. Iida, Y. Anraku, Calcium-sensitive cls mutants of Saccharomyces cerevisiae showing a Pet- phenotype are ascribable to defects of vacuolar membrane H(+)-ATPase activity. J. Biol. Chem. 266,13971-13977 (1991).

43. A. Stathopoulos-Gerontides, J.J. Guo, M.S. Cyert, Yeast Calcineurin Regulates Nuclear Localization of the Crz1p Transcription Factor Through Dephosphorylation. Genes Dev. 13, 798-803 (1999).

**Supplemental Figure Legend**

**Fig. S1. SDS induces local plasma membrane and cell wall damage in budding yeast**

**(A)** Wild type yeast cells were cultured in YPD incubated with or without 0.02% SDS for 3 hr, and then stained with 20 μg/ml calcofluor white for 5 min. The values under the images are % cells with spot signals. The data are presented as the mean±S.D. of four independent experiments. N>100 cells/each experiment. *p*<0.001 by 2-tailed unpaired Student’s *t*-test. Scale bar, 5 μm. **(B)** Wild type yeast cells were cultured at 25˚C under the microscope with 10 μg/ml calcofluor white and then laser damage was induced (yellow stars). Yellow arrows, calcofluor white signal accumulation. The numbers at the upper-left corner indicate time (min). Images were taken at 30 sec intervals. Scale bar, 2 μm. **(C)** Wild type yeast cells expressing Pkc1-GFP or Myo2-GFP were incubated with YPD containing 0.02% SDS for 30 min. The numbers at the lower-left corners indicate % of cells with a polarized GFP signal at the tip of daughter cells. N>200.

**Fig. S2. SDS induces PMD in human cells.**

**(A)** Representative images of DAPI penetration upon 0.008% SDS treatment over time. HeLa cells were cultured in DAPI-containing DMEM with or without 0.008% SDS. Scale bar, 20 μm. **(B)** A representative data set of DAPI influx assay. DAPI-positive cells were counted at indicated time points. N>200. Error bars represents SD. **(C)** Average DAPI intensity of three independent experiments as in (B). Untreated control and 0.008% SDS treatment were significantly different (*p*<0.01) using multiple Welch’s *t*-test with Benjamini, Krieger and Yekutieli correction. **(D)** Flow cytometric analyses of penetrated DAPI signals are shown by histogram overlays. WI-38 cells were cultured in DAPI-containing DMEM with or without SDS (0.0085%). More than 5000 cells were analyzed in each sample. **(E)** Live-cell imaging was performed using HeLa cells cultured in FM1-43-containing DMEM (no FBS) with or without 0.002% SDS. The cytosolic FM1-43 signals were quantified (8 cells per each group). Error bars represents SD.

**Fig. S3. SDS induces local PS externalization on the plasma membrane in human cells; the PS-externalizing spots/blebs colocalize with CHMP4A**

**(A)** A representative data set of Annexin V spots analyses. The number of Annexin V-positive spots per cell was counted. N=60. Error bars represents SD. **(B)** Relative mean Annexin V signals of entire cells were quantified. N>65. **(C)** WI-38 cells were cultured in the medium with 0.008% SDS and pSIVA under the fluorescent microscope. pSIVA binds to externalized PS. Arrow heads, pSIVA positive spots. Yellow arrows, pSIVA positive blebs. The white rectangle region is enlarged in right panels. Scar bar, 20 μm (left), 5 μm (right). Merged images of pSIVA and Bright Field. **(D)** HeLa cells were treated with 0.008% SDS for 1 hr. The cells were incubated with Annexin V-Alexa Fluor 647 conjugate, fixed, and stained with CHMP4A antibody. The white rectangle region in upper panels is enlarged in lower panels. Green: CHMP4A, Red: Annexin V, Alexa Fluor 647 conjugate, BF: Bright Field. White arrows indicate the colocalization of green and red signals. Scale bars, 10 μm (upper), 2 μm (lower).

**Fig. S4. Summary of the screening method**

**(A)** Schematic drawing of the screening method. Blue circle, primary hits (249 mutants); green circle, secondary hits (109 mutants); red circle, confirmed hits (48 mutants). **(B)** Example images of the plates used in the screening.

**Fig. S5. Identification and characterization of factors required for plasma membrane/cell wall damage response in budding yeast**

**(A and B)** Yeast gene deletion mutants were cultured in YPD at 25˚C and then incubated with YPD containing 0.02% SDS. Samples were collected at 30 min intervals and spread onto YPD plates. The number of colonies was counted after 2-3 days. Data are presented as the mean±SD of 3 independent experiments. **(C)** Yeast mutants were treated as in (A) except for the usage of YPD (pH 5.0). **(D)** Yeast mutants were treated as in (A). **(E)** Yeast gene deletion mutants were cultured in YPD at 25˚C and then incubated with YPD containing 0.02% SDS for 30 min. Data are presented as the mean±SD of 3 independent experiments. >300 cells/sample. *p*-values: *, *p*<0.05; **, *p*<0.01; ***, *p*<0.001, compared with wild type+SDS, by 2-tailed unpaired Student’s *t*-test. **(F)** Wild type (Control) and *vma1∆* expressing Crz1-GFP were cultured under the microscope and subjected to laser damage (yellow stars). Images were taken in 30 sec intervals. The numbers at the upper-right corners indicate the time (min). Scale bar, 2 μm. **(G)** Quantification of (F). Data are presented as the mean±SEM of 8 independent experiments. **(H)** Wild type yeast expressing Lact-C2-GFP were cultured under the microscope and subjected to the laser damage as in (F). Relative signal intensity is shown in rainbow pseudocolor. The numbers at the left-top corners indicate the time (min). Arrows: the GFP signal accumulation at the laser damage sites. Scale bar, 2 μm. **(I)** Quantification of (H). Data are presented as the mean of 10 independent experiments. **(J)** Wild type (Control) and *pep3∆* expressing Pkc1-GFP were cultured under the microscope and subjected to laser damage as in (F). Scale bar, 2 μm. **(K)** Quantification of (J). Data are presented as the mean±SEM.

**Fig. S6. Summary of laser damage responses in budding yeast**

Cellular responses after laser damage. Red arrows indicate the laser damage site.

**Fig. S7. SDS induces cellular senescence in human normal fibroblasts**

**(A** **and** **B)** WI-38 cells were treated as in Fig. 3 (D) and (E). SDS concentration was 0.0095%. Scale bar, 20 μm. **(C)** WI-38 cells were treated as in Fig. 3 (G). SDS concentration was 0.0095%.

**Fig. S8. Doxorubicin does not induce plasma membrane damage; KCl treatment induces cellular senescence**

**(A)** Representative images of DAPI penetration upon 0.008% SDS treatment (SDS) and Doxorubicin 250 nM for 24 hr. WI-38 cells were cultured in DAPI-containing DMEM w/ or w/o 0.008% SDS or 250 nM Doxorubicin. Scale bar, 20 μm. **(B)** A representative data set of DAPI influx assay. DAPI-positive cells were counted at indicated time points. N>150. ****; *p*<0.0001 by one-way ANOVA. ns: not significant. Scale bar, 100 μm.

**(C and D)** WI-38 cells were treated with 75mM KCl for 24 hr, washed, and released into fresh medium. SA-β-gal-positive cells were detected using the cells seven days after KCl wash away. Scale bar, 50 μm. Graphs show quantification of SA-β-gal-positive cells (n>100). Data are presented as the mean±SEM of 3 independent experiments. *p*-value, by 2-tailed unpaired Student’s *t*-test. **(E)** WI-38 cells were treated with 75mM KCl for 24 hr, washed, and released into fresh medium. Samples were collected at indicated time points. Western blot was performed as in Fig. 3(G).

**Supplemental Table**

**Table S1. Summary of GO enrichment analysis results.**

**Table S2. Summary of *mod*PhEA analysis results.**

**Table S3. 18 out of 48 screening hits are reported to have altered replicative lifespan.**

**Table S4. Yeast strains used in this study.**

**Table S5. Antibodies used in this study.**

**Table S6. Primer sequences used for qPCR analyses.**
